# A Cyclisation and Docking Protocol for Cyclic Peptide-Protein Modelling using HADDOCK2.4

**DOI:** 10.1101/2022.01.21.477251

**Authors:** Vicky Charitou, Siri C. van Keulen, Alexandre M.J.J. Bonvin

## Abstract

An emerging class of therapeutic molecules are cyclic peptides with over 40 cyclic peptide drugs currently in clinical use. Their mode of action is, however, not fully understood, impeding rational drug design. Computational techniques could positively impact their design but modeling them and their interactions remains challenging due to their cyclic nature and their flexibility. This study presents a step-by-step protocol for generating cyclic peptide conformations and docking them to their protein target using HADDOCK2.4. A dataset of 30 cyclic peptide-protein complexes was used to optimize both cyclisation and docking protocols. It supports peptides cyclized via an N- and C-terminus peptide bond and/or a disulfide bond. An ensemble of cyclic peptide conformations is then used in HADDOCK to dock them onto their target protein using knowledge of the binding site on the protein side to drive the modeling. The presented protocol predicts at least one acceptable model according to CAPRI criteria (Critical Assessment of Prediction of Interaction) for each complex of the dataset when the top 10 HADDOCK-ranked single structures are considered (100% success rate top10) both in the bound and unbound docking scenario. Moreover, its performance in both bound and fully unbound docking, is similar to the state-of-the-art software in the field, Autodock CrankPep. The presented cyclisation and docking protocol should make HADDOCK a valuable tool for rational cyclic peptide-based drug design and high-throughput screening.

## Introduction

Cyclic peptides are promising therapeutic molecules with about one new cyclic peptide drug entering the market every year^1^. Their success lies in their favorable pharmacokinetic characteristics, such as their enhanced metabolic stability^2^ and oral availability. Cyclic peptides are usually more stable than their linear counterparts which is mainly due to resistance to chemical or enzymatic hydrolysis^3^. Furthermore, cyclic peptides can bind large, flat protein surfaces with high affinity and specificity^4^ as well as disrupt protein-protein interactions^5–8^. Cyclic peptides are therefore considered a promising compound class for therapeutic modulation of challenging protein-protein interactions.

Examples of cyclic peptides in clinical use are the immunosuppressant drug cyclosporin A^9^, antibiotics such as vancomycin^10^ and gramicidin S^11^ as well as antifungals^12^. Despite their successful applications, the mode of action of the majority of these molecules is poorly understood^13^. Currently, the optimization of cyclic peptides is mainly an empirical pursuit^14–16^ involving the synthesis of many different analogs in the hope of finding one with improved target-binding properties while often facing significant delays and synthetic challenges^13,17–19^. New high-throughput screening approaches for macrocyclic peptides, such as RaPID^16^, attempt to overcome these challenges. However, these methods are limited to some types of cyclic peptides and require further optimization. In these cases, computational techniques can often complement experimental work, as shown by Goldbach *et al*^20^. In that study, RaPID^16^ peptide selection in combination with HADDOCK was used to predict protein-peptide complexes whose complex form was challenging to solve experimentally, demonstrating cyclic peptide discovery could greatly benefit from computational techniques.

Similar to linear peptides, cyclic peptides present computational challenges as they include a large number of rotatable bonds, often lack secondary structural elements and their conformational transitions might be difficult to sample due to their cyclic nature. All these aspects pose challenges for predicting protein-peptide complexes^21,22^. To overcome such challenges, a common approach is to generate cyclic peptide conformational ensembles prior to docking. These can be obtained via methods such as Monte Carlo as well as molecular dynamics (MD) simulations (e.g. high-temperature MD simulations and replica-exchange MD)^23^. The generated ensemble can then be used as input to model the protein-peptide complexes. Apart from difficulties raised due to the need of generating initial cyclic conformations, these procedures can also be time-consuming and require expertise on various software packages.

To overcome this issue and offer users a single software solution, two docking packages, AutoDock^24^ and Glide^25^, introduced a macrocycle module. Using docking instead of Monte Carlo or MD simulations for the prediction of cyclic peptide-protein complexes offers an increase in efficiency, enabling high-throughput screening of different peptides. However, peptide-protein docking remains challenging as it is difficult to incorporate the conformational sampling of such flexible peptides in docking calculations. Currently, the state-of-the-art practice for cyclic peptide docking is AutoDock CrankPep (ADCP)^26,27^ that offers a one-software pipeline to generate a cyclic peptide conformational ensemble and performs docking with the obtained cyclic peptide models. ADCP folds the peptide in the energy landscape created by the receptor, thus yielding docked peptide poses.

In this study, a new cyclic peptide cyclisation and docking protocol is presented using the integrative modelling software package HADDOCK^28^. Nine different docking protocols were benchmarked on a set of 30 cyclic peptide-protein complexes to evaluate their performance. The best performing protocol shows that HADDOCK achieves a competitive performance in the field for both bound (*holo*) and unbound (*apo*) receptor conformations while using binding interface information on the receptor side in combination with peptide conformations generated from sequence. More specifically, HADDOCK predicts within the top 10 HADDOCK-scored solutions of bound (*holo*) receptor docking, at least one medium or higher quality structure for 70% of the tested cyclic peptide-protein complexes, according to the CAPRI^29^ (Critical Assessment of Prediction of Interaction) criteria. In unbound (*apo*)receptor docking, HADDOCK predicts models in the same quality range within the top 10 solutions for 60% of the dataset which can be further enhanced by clustering. When only short cyclic peptides (≤10 residues) are considered, the performance of HADDOCK further increases, reaching 88.2% in bound receptor docking and 68.75% for fully unbound docking. A success rate of 100% is reached for both holo and apo receptor conditions when considering acceptable or higher quality models for the short cyclic peptide subset. Overall, HADDOCK’s performance is comparable or better than the current state-of-the-art practice, especially for short cyclic peptides.

## Methods

### Dataset Preparation

The two datasets used to benchmark the protocols in this study are composed of 30 cyclic peptide-protein complexes extracted from the dataset described by Zhang et al^26^. The first set of complexes, the Backbone dataset, includes 18 complexes in which the peptide is cyclized through its N- and C-termini with a minimum sequence length of six amino acids (Table S1). Four out of these 18 peptides include more than 10 residues and contain an additional disulfide bond (Figure 1). The second set of complexes, the Disulfide dataset, includes 12 complexes which contain peptides cyclized through a single disulfide bond (Table S2).

**Figure 1:**
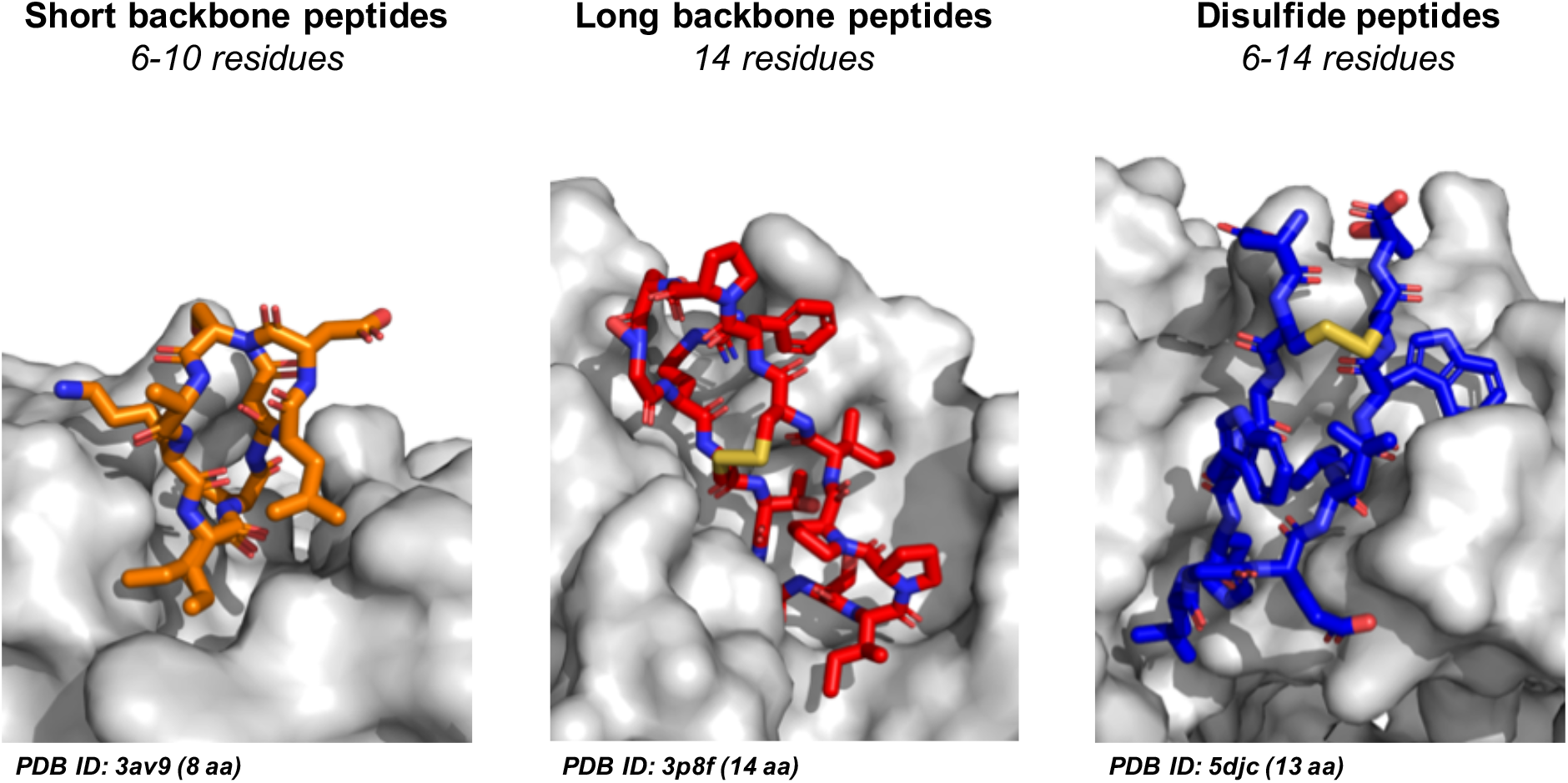
Visualization of Backbone and Disulfide cyclic peptide datasets. Stick representation of 3 peptide examples from the Backbone and Disulfide dataset indicating their PDB ID and sequence length. Receptor proteins are shown in a gray surface representation and the peptides are highlighted in orange, red and blue. Short Backbone peptides (orange) consist of maximum 10 residues whereas long Backbone peptides (red) can include up to 14 residues and an additional disulfide bond. Disulfide peptides (blue) range in length between 6 and 14 residues.

All cyclic peptide-protein complexes in both the Backbone and Disulfide datasets were selected based on the following criteria: (i) length of the peptide (from 6 up to 14 residues) and (ii) formation of only one disulfide bond. In case of the Disulfide dataset, an additional criterium of no more than 2 residues that would not be taking part in the cyclization was imposed as well to exclude peptides with too disordered termini (Figure 1, Table S2).

The bound structure of each receptor in the two datasets was extracted from the corresponding Protein Data Bank (PDB)^30^ entry of the complex. For 25 out of 30 receptors an unbound form (apo) was available (Table S1 and S2). The protein structures were prepared for HADDOCK calculations via pdb-tools^31^. The holo protein structures were isolated from the available complex using `pdb_splitchain` and chain ID information was removed with `pdb_chain`. Hetero atoms were removed with `pdb_delhetatm`. A similar approach was employed for the preparation of the unbound (apo) receptor structures. In case the receptor was a multi-chain protein, extra steps were needed to properly prepare the PDB file^32^. Since the molecule will be assigned a single chain ID for docking in HADDOCK, it is important that there is no overlap in the residue numbering between the chains. To ensure that, `pdb_reres` was used from pdb-tools^31^. However, if gaps are present in the sequence of the protein, a more appropriate alternative is the `pdb_shiftres` script to ensure gap preservation. Furthermore, in case the receptor structure contained missing loops and/or gaps, distance restraints were defined to keep the different chains together during the high temperature flexible refinement stage of HADDOCK. These were generated with the `restrain_bodies.py` script from haddock-tools^33^. The output file was saved as `hbonds.tbl` and was used by HADDOCK as hydrogen bond restraints, by activating the hbond setting (Table S6). Finally, conformations of the cyclic peptides were generated using an optimized cyclisation protocol through a combination of PyMOL^34^ and HADDOCK (see below).

### Peptide Cyclisation Protocol

The cyclic peptides used for peptide-protein docking were prepared with PyMOL^34^ and the integrative modelling software package HADDOCK^28^, version 2.4, starting from the available peptide sequences. The cyclisation protocol consists of three steps (Figure 2):

1. Generating the starting conformations from the peptide sequence (PyMOL)
2. Reducing the distance between the termini/disulfide-bond of the peptide (HADDOCK)
3. Cyclizing the peptide (HADDOCK)

**Figure 2:**
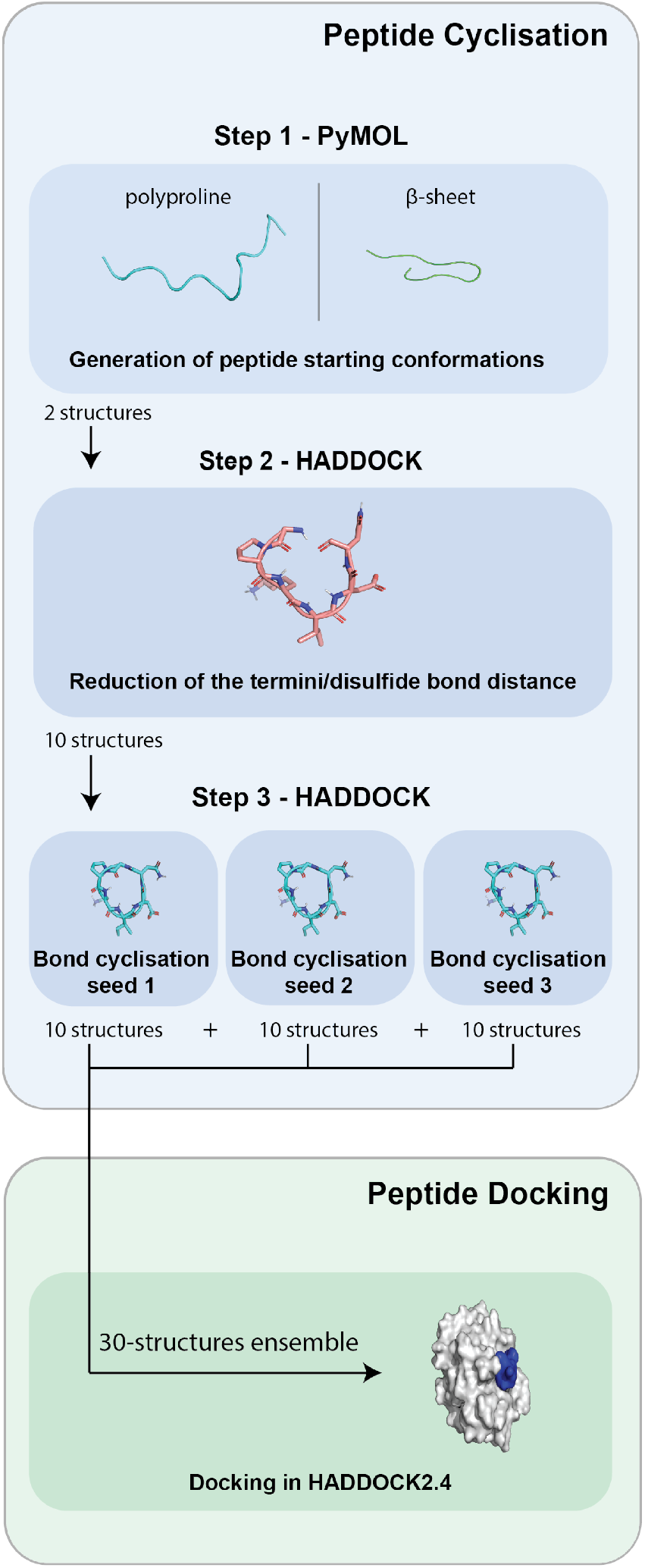
Flow chart of the peptide cyclisation and docking protocol. In Step 1, the starting conformations (polyproline and β-sheet) are generated from the peptide’s sequence. These two starting conformations are the input for Step 2 in which the distance between the termini/disulfide bond of the peptide is reduced by applying distance restraints. 10 representative structures are used as an input for Step 3. In Step 3, the cyclisation bond is formed and by repeating Step 3 with different seeds the ensemble of peptide conformations is created. This ensemble is later used for docking the cyclic peptides with their respective receptors.

#### Step 1: Generating the Peptide Starting Conformations

Using the PyMOL utility script `build_seq`, written by Robert Cambell^35^, each peptide was generated in a beta-sheet (`ss=beta`) and a polyproline (`ss=polypro`) conformation (Figure 2). Two peptides from the disulfide dataset were crystalized with capped termini, namely an N-terminal acetyl group (ACE) and a C-terminal N-methyl amide group (NME). For these peptides (PDB ID: 3wnf and 5h5q) a glycine residue was added to both termini which was then mutated using the pdb-tools script `pdb_mutate` to an ACE or NME cap. Uncapped N- and C-termini in the disulfide dataset were kept positively and negatively charged by introducing an NH3^+^ group at the N-terminus and a negatively charged carboxylate moiety on the C-terminal side. All the other, nonterminal residues were protonated according to physiological pH levels. Subsequently, the PyMOL-generated structures were used as an input for the first cyclisation step in HADDOCK, Step 2.

#### Step 2: Reducing the Distance Between the Termini/Disulfide Bond of the Peptide

The two starting conformations from Step 1, the beta-sheet and polyproline state, were used as an ensemble of peptide conformations for the HADDOCK2.4 calculations in Step 2 to generate models with a reduced distance between their termini/disulfide bond. This is a preparatory step before the actual cyclisation of the peptide (introduction of the covalent bond) which is performed in Step 3 (Figure 2).

The main use of HADDOCK is to perform biomolecular docking of N≥2 molecules, but here the software package is first used in single molecule mode in the cyclisation protocol to generate the cyclized peptides. The default HADDOCK protocol includes three stages^28^. The first stage is rigid body docking (it0) and is usually guided by experimental or predicted information about the protein-protein interface introduced as distance restraints. The best HADDOCK-scored models (default: 200) from it0 continue to the next stage (it1), which is a semi flexible refinement in torsion-angle space. In it1, three simulated annealing refinements are performed in which flexibility is gradually introduced into the system. After a high temperature rigid body search (default: 500 steps), the first cooling simulated annealing step (default: 500 steps) is performed in which the proteins are treated as rigid bodies and their respective orientation is optimized. In the second simulated annealing step (default: 1000 steps), the side chains at the interface are allowed to move. In the third simulated annealing step (default: 1000 steps), both side chains and backbone at the interface can move to allow for conformational rearrangements. All models from it1 move to the final refinement stage (itw) in which the protein-protein interface is refined by either an energy minimization (default in HADDOCK2.4) or by a short molecular dynamics simulation in explicit solvent.

For the cyclisation of the peptides (Step 2 and Step 3) HADDOCK was run for a single molecule with the default docking protocol modified as follows (Table S3 and S4):

- In it0, it1 and itw, the number of structures generated was set to 400
- In it0, it1 and itw, the cyclic peptide was defined as fully flexible
- In it0, 400 initial structures were generated while the rigid body stage was skipped
- In it1, all steps were increased by a factor of 4
- In itw, the final refinement was performed with an explicit water shell
- The generated models were clustered using pairwise root-mean square deviation (RMSD)^36,37^ with a cut-off of 2.5 Å.

In Step 2, the electrostatic energy term is also turned off for the semi-flexible simulated annealing (it1) stage (Table S3). This is because with electrostatics switched on, the generated peptide conformations tend to have their charged/hydrophilic side chains pointed towards the inside of the peptide cyclic center, leading to an unphysical conformation of the cyclic peptide which is further away from the holo peptide state. In addition, distance restraints are introduced between the peptide’s termini/residues forming the disulfide bonds to reduce the distance between the atoms of the peptide that are involved in bond formation:

- *Disulfide dataset*: The distance restraints used were between the Cα-Cα atoms of the cysteine residues that form the disulfide bond and defined with a 4 Å upper limit. Likewise, the distance between the Cβ-Cβ atoms was defined as 3.5 Å and the distance of the Sγ-Sγ atoms was defined as 2 Å. Both upper-bound and lower-bound corrections were 0.1 Å (Figure 3A). The Cα-Cα restraint was used to bring the cysteine residues closer together and the Cβ-Cβ distances were selected to restrain the angle of the side chains of the cysteines.
- *Backbone dataset*: Backbone distance restraints, C-N of 1.3 Å and O-N of 2.3 Å, were used with an upperbound and lower-bound correction of 0.1 Å (Figure 3B). The O-N distance is defined to restrain the O-C-N angle. When peptides are cyclized through their termini and include an additional disulfide bond (PDB ID: 1smf, 4k1e, 4kel, 3p8f), all aforementioned disulfide and backbone distance restraints (termini [C-N : 1.3 Å, ON : 2.3 Å], disulfide [Cα-Cα : 4 Å, Sγ-Sγ : 2 Å]) were used except for the Cβ-Cβ restraint, which proved not to be required in this case (Figure 3C).

**Figure 3:**
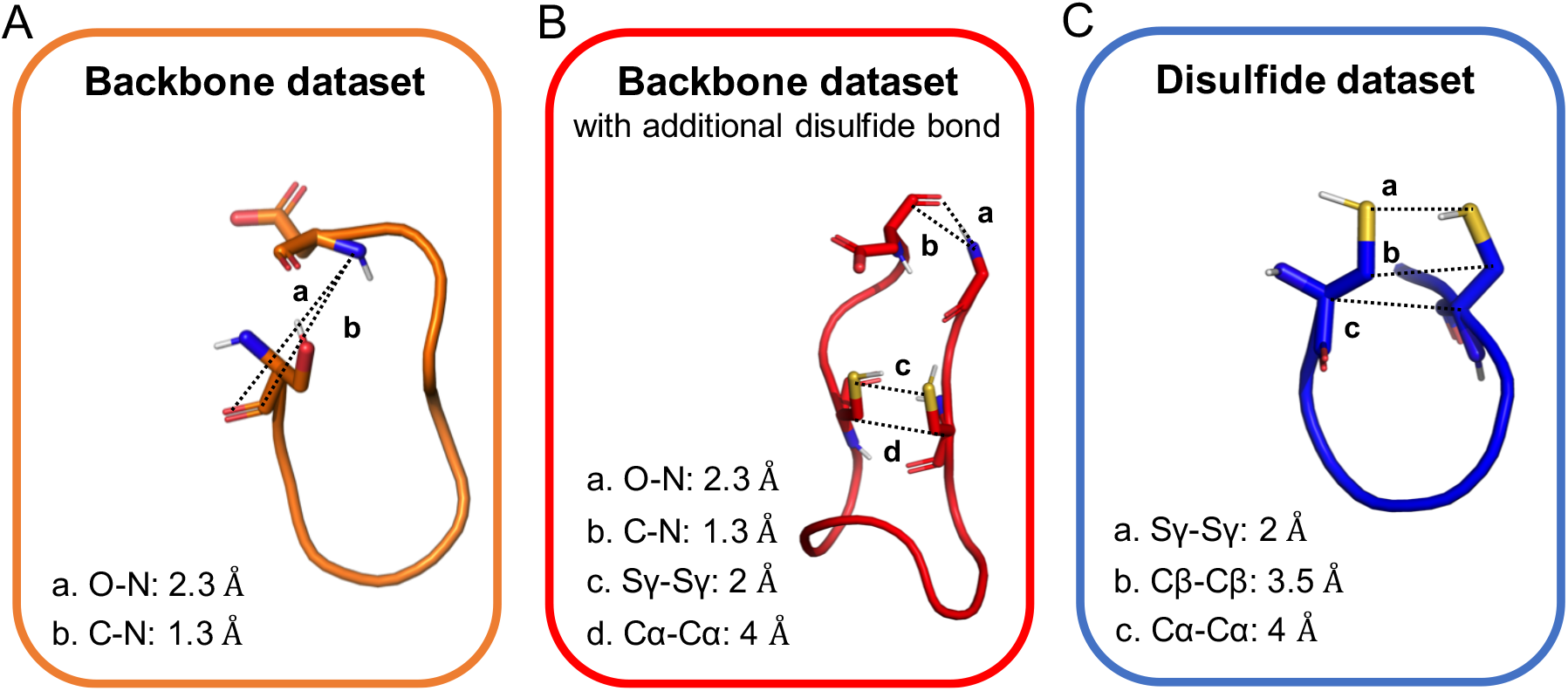
Visualization of distance restraints defined in the cyclisation protocol. (A) Backbone restraints for short peptides (≤10 residues). (B) Backbone restraints for long Backbone peptides including an example of an additional disulfide bond. (C) Disulfide restraints for peptides cyclized through a disulfide bond only.

#### Step 3: Forming the Covalent, Cyclic Bond of the Peptide

From the modelled peptides of Step 2, one representative model was selected from each of the 10 most populated clusters after RMSD clustering (Figure 2). The center of each selected cluster was used as a representative model. These ten representative models composed the ensemble of peptide conformations used in Step 3 of the cyclisation protocol.

Similar to Step 2, the electrostatic energy term was turned off in it0 and it1 during cyclization in Step 3. The same unambiguous distance restraints used in Step 2 for the termini/disulfide bonds were included in Step 3 as well. In addition, the cyclic peptide option in HADDOCK is set to true for the covalent bond to be defined (provided the N and C atoms are within 3.5 Å distance (increased from the default 2Å)). The disulfide bond is automatically recognized and defined provided the S-S distance is ≤4A (increased from the default 3Å) (Table S5).

The difference in Step 3 compared to Step 2 is that the restraints are only implemented in the first two stages of docking (it0 and it1). The final stage, itw, is an extended water refinement with distance restraints switched off and water steps increased by a factor of 2 (Table S4). Hence, the final itw stage is an unrestrained very short MD refinement of the peptide models.

#### Generating the Ensemble of Cyclic Peptide Conformations with Step 3 for Peptide-Protein Docking

From the third and final step in the peptide cyclisation protocol, a maximum of ten representative models are selected out of the 10 most populated clusters after RMSD clustering. If less than 10 clusters are formed during Step 3, only the representative structures from the available clusters are used. The selected models correspond to the center of the cluster – called representative models in the following. The ensemble of cyclic peptide conformations obtained from Step 3 is subsequently used for peptide-protein docking with the bound (*holo*) or unbound (*apo*) form of the receptor in HADDOCK (see below). The size of the cyclic peptide ensemble can be increased by repeating Step 3 with a different initial HADDOCK seed, which is used by the CNS random number generator to produce different initial velocities for the peptide. For example, by repeating Step 3 three times with a different seed, a peptide ensemble with a maximum of 30 conformations can be generated for each peptide (Figure 2). By increasing the number of cyclic peptides in the ensemble, a larger variation in peptide conformation can be included during docking.

### Cyclic Peptide-Protein Docking Protocol

After the peptide cyclisation process, the obtained cyclic peptide ensemble and the respective receptor structure are used to perform peptide-protein docking with HADDOCK2.4. Nine different docking protocols were tested in order to find the one leading to the best quality of docked models (Table 1). These protocols differ in the size of the cyclic peptide ensemble, the flexibility treatment and the presence/absence of a final refinement in explicit solvent in itw. The docking protocols were benchmarked on the Backbone dataset using the *holo* (bound) structure of their receptor. The Backbone dataset was chosen for this as its size is larger than the Disulfide peptide dataset. Protocol benchmarking was done using the *holo* receptor rather than the *apo* structure to concentrate on the peptide part of the docking setup and to evaluate the effect of the various settings.

**Table 1:** Overview of the tested peptide-protein docking protocols.

| Protocol | Cyclic Peptide Ensemble <sup>a</sup> | Extra Flexibility<sup>b</sup> | Explicit Solvent Shell<sup>c</sup> | ID |
| --- | --- | --- | --- | --- |
| 1 | 5 | OFF | OFF | 5STR |
| 2 | 30 | OFF | OFF | 30STR |
| 3 | 40 | OFF | OFF | 40STR |
| 4 | 50 | OFF | OFF | 50STR |
| 5 | 60 | OFF | OFF | 60STR |
| 6 | 50 | ON | OFF | 50STR_FLEX |
| 7 | 50 | OFF | ON | 50STR_SOLVSHELL |
| 8 | 50 | ON | ON | 50STR_FLEX_SOLVSHELL |
| 9 | 50 | ON/OFF<sup>d</sup> | ON/OFF<sup>d</sup> | 50STR_COMB |

a. Maximum number of conformations in the cyclic peptide ensemble
b. Parameters changed in `run.cns` for peptide flexibility when peptide is defined as molecule 2: `nfle_2` (changed from 0 to 1), `start_fle_2_1` and `end_fle_2_1` (changed to the residue numbering of the first and last residue, respectively).
c. Parameters changed in `run.cns` file for the itw stage: `solvshell` (changed from false to true)
d. Extra flexibility and explicit solvent is switched ON for long cyclic peptides (>10 residues) and switched OFF for short cyclic peptides ( $\leq 10$ residues)

All nine protocols include the same definition of the interface on the receptor side which is used by HADDOCK to guide the docking process. Ambiguous interaction restraints (AIRs) describing the interaction region were used to guide the docking in which the peptide was defined as fully passive and the receptor binding region as active. The protein interface was defined by taking into account all receptor residues within a 5 Å distance cutoff of the cyclic peptide in the available holo complex structure. By defining the peptide as fully passive, regions of the peptide that do not contact the protein will not be penalized. This AIR definition is in line with the best practice guide for peptide-protein docking with HADDOCK^38^. The AIRs file that was used during the docking calculations was generated by using the `active-passive-to-ambig` script from haddock-tools^33^.

The base docking setup in all nine protocols, includes the following parameters changes with respect to the default settings (Table S6):

- The peptide was defined as cyclic by activating the cyclic peptide option in `run.cns`
- In it0, it1 and itw, the number of models docked were increased to 5000, 400 and 400, respectively
- In it1, all steps in the semi-flexible simulated annealing stage were increased by a factor of 4 to allow for more conformational sampling of the peptide in the context of the receptor
- Generated models were clustered based on the interface ligand RMSD using a 5 Å cutoff

#### Assessing the Impact of the Peptide Ensemble Size (protocols 1-5)

In the first five protocols, the maximum number of conformations in the cyclic peptide ensemble were varied from 5 to 60. Protocol 1 includes the smallest ensemble of peptide conformations of 5 representatives selected from the top 5 most populated clusters, generated using a single seed. In protocol 2-5, the size of the ensemble was increased by using 3, 4, 5 or 6 different seeds and selecting a maximum number of 10 representatives from each seed, depending on the number of available clusters. The total number of representatives used per protocol are shown in Tables S7 and S8.

#### Assessing the Impact of Allowing for Extra Flexibility and Use of Explicit Solvent Shell (protocols 6-9)

In protocols 6-9, the effect of extra flexibility and explicit solvent refinement was assessed. Because of the intrinsically high flexibility of peptides, the optimal setting for protein-peptide docking of linear peptides is to define the peptides as fully flexible and to refine them in an explicit solvent shell. By defining a peptide as fully flexible, both the side chains and the backbone of the entire peptide are allowed to move throughout the docking, except for the initial rigid body minimization stage (it0). To test whether this setting for linear peptides is also required for cyclic ones, the peptides were defined as fully flexible throughout stage it1 and itw in protocol 6. On the other hand, to evaluate the impact of explicit solvent shell refinement in cyclic peptides, protocol 7 was used, in which the peptides were refined in explicit solvent (water) but the default flexibility was maintained. Moreover, to test the combined effect of these two settings in protocol 8, peptides were both defined as fully flexible and refined in an explicit solvent shell. Finally, in protocol 9, peptides were treated differently based on the length of their sequence. For peptides that are composed of 10 residues or less, the default flexibility was applied and they were only subjected to a final energy minimization. Peptides longer than 10 residues were treated as fully flexible throughout it1 and itw and refined in an explicit solvent shell.

### Success-Rate Analysis based on the Fraction of Native Contacts

The quality of the generated models was assessed using the fraction of native contacts (f_nat_). This metric is defined as the number of native (true) residue-residue contacts in the predicted complex at the protein-peptide interface, divided by the number of contacts in the reference crystal structure of the complex^29^. A pair of residues on different sides of the protein-protein interface was considered to be in contact if any of their heavy atoms are within 5 Å distance cutoff. According to the CAPRI^39^ criteria for protein-peptide complexes, models with an f_nat_ above 0.2 were ranked as acceptable, above 0.5 as medium and above 0.8 as high quality. The interface root-mean square deviation (i-RMSD) is another metric typically used for interface accuracy. However, the i-RMSD is less well suited for cyclic peptide analysis, especially for cases in which flexible terminal extensions exist outside the cyclic structure (i.e. for the disulfide dataset)^26^, which is discussed in the Supplementary Methods section and Figure S1. Nevertheless, i-RMSD values were also calculated for the generated models along with f_nat_ and are reported in the supplementary information (see Table S11). According to CAPRI^39^ criteria for protein-peptide complexes, models with i-RMSD values below 2.0 were ranked as acceptable, below 1.0 as medium and below 0.5 as high quality.

The performance of each tested protocol was evaluated using the f_nat_ success rate. The success rate for single structure analysis is defined as the percentage of cases in which at least one model of a given accuracy (high, medium or acceptable) is found within the top *N* solutions ranked by HADDOCK (*N*= 1, 5, 10, 20, 50, 100, 200) using the itw scoring function^40^. For example, a success rate of 60% for medium quality structures in a top 10 means that in the top 10 HADDOCK-ranked solutions at least one model of medium or higher quality was found for 60% of the complexes of the dataset. Regarding the cluster analysis, the success rate is calculated for the four best clusters, according to the itw HADDOCK score, using the top 4 structures per cluster. The cluster success rate is defined as the percentage of cases in which at least one model of a given accuracy (high, medium or acceptable) within the top 4 members of the cluster is found within the best N clusters ranked by the itw HADDOCK score (N= 1,2,3,4) and considering.

## Results and Discussion

As the quality of the generated cyclic peptide structures can greatly influence the peptide-protein docking performance, the quality of the cyclic peptide ensembles generated with HADDOCK are first discussed. Subsequently, the different docking protocols are compared to determine the optimal settings for cyclic peptide docking. These have been tested using the Backbone dataset and the *holo* structure of the receptor. The best cyclic peptide docking protocol was then applied to fully unbound docking using the *apo* receptor structure, for both the Backbone and Disulfide datasets. Next, the optimized protocol’s performance is compared to the baseline docking performance for both datasets, using the holo receptor and single holo cyclic peptide conformation. Finally, the HADDOCK docking results are compared to the results of cyclic peptide docking using Autodock CrankPep.

## Quality Assessment of the Cyclic Peptide Conformational Ensemble

Prior to docking, the quality of the generated cyclic peptide ensembles was analyzed by calculating their backbone RMSDs with respect to the holo cyclic peptide conformation.

### The Peptide Ensemble of the Backbone Dataset is Closer to the Holo State than the Disulfide Dataset

When considering the best RMSDs obtained within the tested ensemble sizes (from 5 to a maximum of 60 structures), the average RMSD value of the best structures becomes lower by enlarging the ensemble (Tables S9 and S10). This trend suggests that by enlarging the peptide conformational ensemble, the probability of including a peptide structure resembling the holo peptide conformation is increased. The presence of a near holo peptide conformation in a peptide ensemble could potentially lead to enhanced docking results. The RMSD analysis of the 50-structures peptide ensemble shows that the modelled peptides from the Backbone dataset are closer to their *holo* crystal structure than the cyclic disulfide ones (Figure 4A). On average, the best generated Backbone peptides of each ensemble have an RMSD of 1.5-2 Å, while the best Disulfide peptides only reach an RMSD of 3 Å (Table S9, S10).

**Figure 4:**
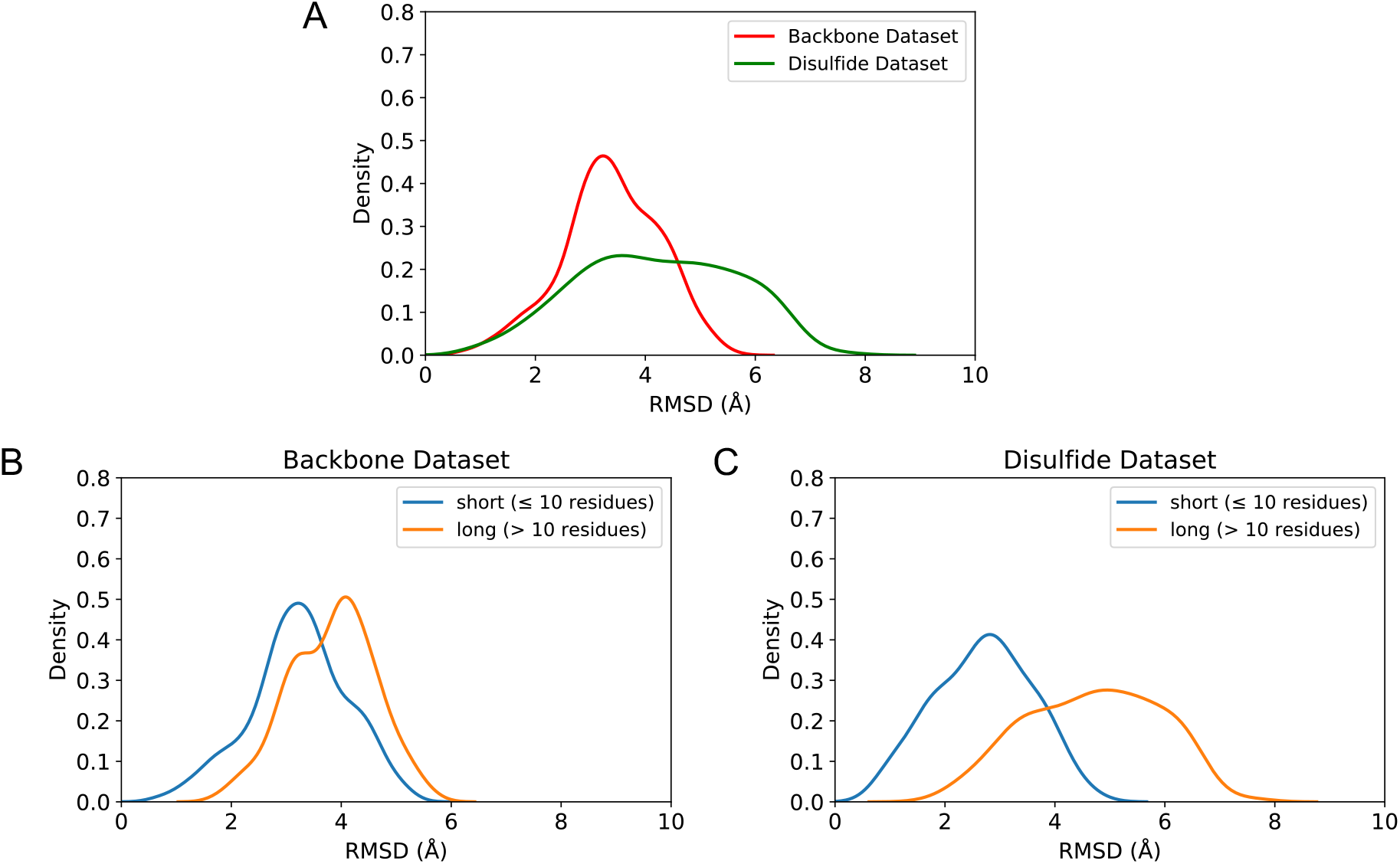
RMSD distribution of the 50-structure cyclic peptide ensemble. (A) The backbone RMSD distribution of the generated Backbone peptides, depicted in red, calculated by using 835 data points (18 peptides*representative structures/seed), while 416 data points were included for the Disulfide dataset shown in green (12 peptides*representative structures/seed). The number of representative structures per seed varies between 1 to 10 for every peptide and depends on the number of clusters formed at the end of the cyclisation step (Step 3) (Table S7, S9). For PDB entries 2ck0, 4ou3, 4ib5, 5h5q, 5djc and 5wxr of the Disulfide dataset, the residues outside their cyclic region were excluded during the RMSD calculation. (B) The backbone RMSD distribution of the short (blue) and long (orange) Backbone peptides was calculated using 635 (14 peptides*representative structures/seed) and 200 (4 peptides*representative structures/seed) data points, respectively (Table S7). (C) The backbone RMSD distribution of the short (blue) and long (orange) Disulfide peptides was calculated using 98 (3 peptides*representative structures/seed) and 318 (9 peptides*representative structures/seed) data points, respectively (Table S8). For PDB entries 2ck0, 4ou3, 4ib5, 5h5q, 5djc and 5wxr of the Disulfide dataset, the residues outside their cyclic region were excluded during the RMSD calculation.

A reason for the difference in ensemble quality between the two datasets could be the flexible termini of the Disulfide peptides. However, in the RMSD calculations of the Disulfide dataset only the cyclic regions of the peptides were considered to eliminate differences caused by disordered termini. Therefore, the impact of peptide length was assessed next as extending the peptide length increases the number of degrees of freedom, leading potentially to a higher backbone RMSD.

### The Conformational Quality of the Short Cyclic Peptides is Higher than the Quality of the Long Peptides in Both Datasets

Regarding the quality of the generated cyclic peptides separated by sequence length, the shorter peptides (≤ 10 residues) are closer to their *holo* crystal structure (Figure 4B, 4C) than the long cyclic peptides. The Backbone dataset consists of mainly short (14 out of 18) peptides (Table S1), which could explain the overall higher quality of the conformational ensemble compared to the Disulfide peptides which only contains 3 short peptides out of 12 (Figure 4A) (Table S2). This difference in structural quality between short and long peptides can be observed in both the Backbone and the Disulfide datasets and points to the challenges imposed by *ab-initio* modelling when the number of rotatable bonds increase^41^.

## Cyclic Peptide-Protein Docking Protocol Optimization

After the quality assessment of the generated peptide ensembles, optimization of the cyclic peptide-protein docking protocol was performed. Nine protocols were tested to determine the best settings for cyclic peptide docking using HADDOCK2.4. The performance of the different protocols was assessed by using f_nat_ as a metric for the success-rate analysis (see Methods). Both single structure and cluster analyses were performed for the best performing protocol (Figure S2).

The inherent flexible nature of peptides poses difficulties in docking due to the possible conformational changes occurring upon binding. Since cyclic peptides are a sub-category of peptides, the search for the optimal docking settings was based on the previously proposed HADDOCK protocol for linear peptide-protein docking^42,43^, a protocol that attempts to overcome aforementioned difficulties by including the following features:

- An ensemble of three peptide conformations (beta, polyproline-II and alpha helical) is used for docking
- In it1 and itw, the peptides are defined as fully flexible
- In itw, the models are refined via a short MD simulation in an explicit solvent shell (the default in HADDOCK2.2)

For cyclic peptide-protein docking, the effect of these three settings was evaluated by performing the protocols described in Table 1. First, the size of the peptide conformational ensemble was assessed by comparing the results of protocols 1-5 (Table 1) in which the peptide ensemble size increases from 5 to 60 conformations. Second, protocols 6-9 (Table 1) were conducted to determine the effect of extra flexibility and explicit solvent shell refinement.

### 50 Peptide Structures is the Optimal Ensemble Size for Cyclic Peptide Docking

To identify the optimal number of structures per ensemble, experiments of various ensemble sizes were performed. Ensembles of 5, 30, 40, 50 and 60 structures (5STR, 30STR, 40STR, 50STR, 60STR) were generated per peptide included in the Backbone dataset (see Methods). In order to concentrate on the peptide conformational sampling, the docking was performed with their respective *holo* receptor structure (Table 1, Table S5). The results in Figure 5 show that increasing the number of structures of the ensemble from 5 to 30 leads to an increase of 11.1% in the success rate of medium or high-quality structures within the top 10 HADDOCK-scored solutions. A further expansion of the ensemble to 50 structures resulted in complex models that continued showing this ascending trend in performance. The medium or higher-quality success rates with a 50-structures ensemble were 55.5% for the top 5 HADDOCK-ranked solutions and 61.1% for the top 10 (Figure 5).

**Figure 5:**
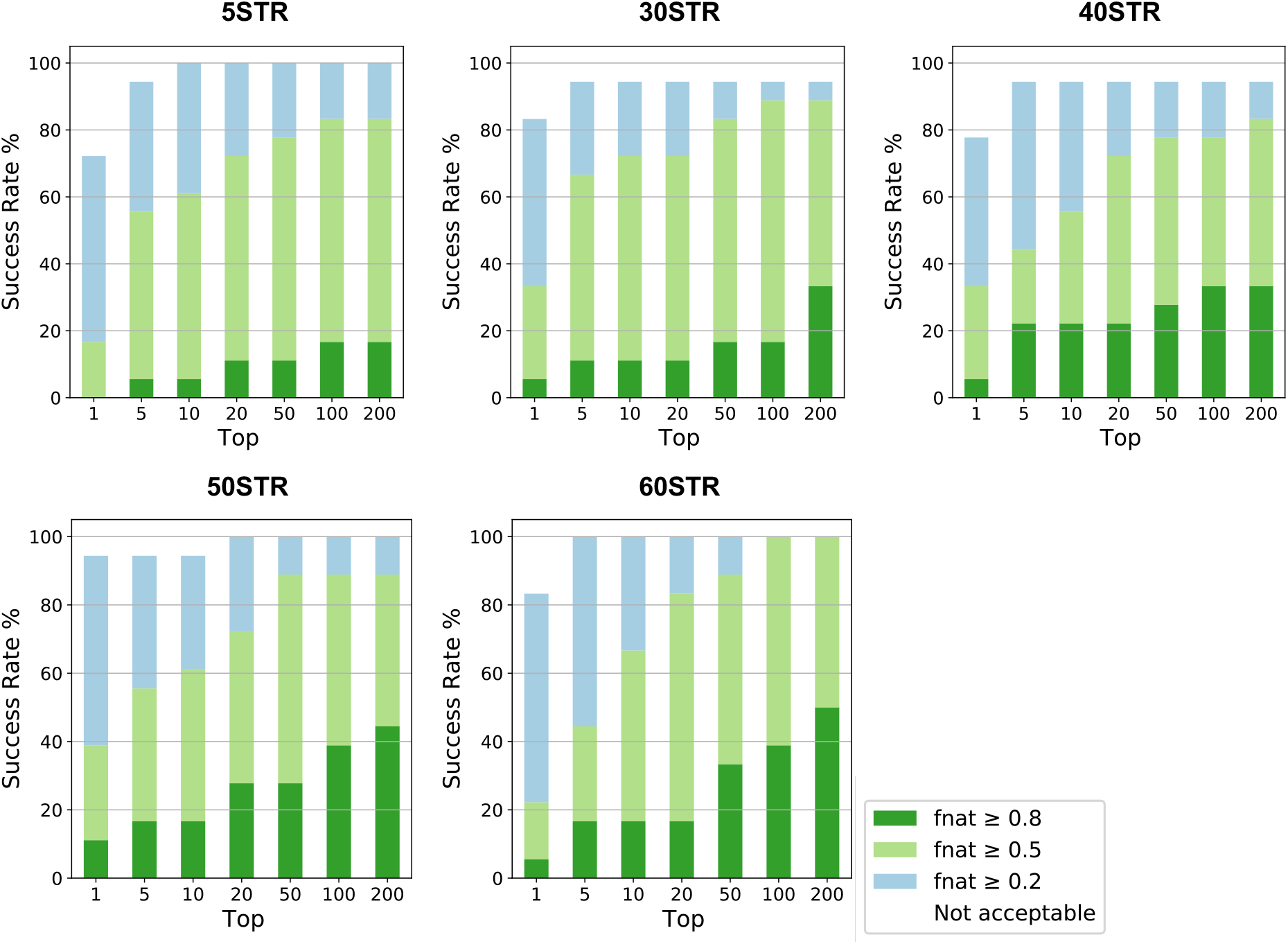
F_nat_ success-rate plots for 5STR, 30STR, 40STR, 50STR and 60STR docking protocols. Plotted are the success rates (%) of protocols 1-5, including a 5, 30, 40, 50 or 60-structures ensemble. The color coding (blue, light green, green) indicates the quality of the models, from acceptable to medium and high quality, according to CAPRI criteria.

However, for the success rates of the 60-structures ensemble experiment, a reduction in quality can be observed in comparison to the 50-structures setup (Figure 5). A reason for this reduction could be the “dilution effect”, which is caused by enlarging the peptide ensemble without increasing the number of models generated in the initial rigid body docking (it0, 5000), leading to a decrease in in sampling per conformation during docking. However, after increasing the number of generated it0 models to 10 000 in the 60-structures setup, no significant increase in performance was observed (Figure S3). Another reason for the reduction in model quality could be the scoring of the generated models: By adding more conformations to the ensemble, challenges could be introduced to the scoring of the docked models, leading to more false positive solutions to be ranked higher in the top. This is illustrated in Figure 5: Although the 60STR experiment generates more medium and high-quality structures in the top 200, the scoring function fails to rank them within the top 10. Re-optimization of the scoring function for cyclic peptides could potentially improve the scoring.

### Extra Flexibility and Explicit Solvent Refinement are Required Only for Long Peptides (>10 residues)

After identifying the best performing ensemble size, the effect of extra flexibility and explicit solvent shell refinement during cyclic peptide docking was evaluated. In our previous work on linear peptide structures, the peptides were treated as fully flexible and refined in an explicit solvent shell ^43^. To evaluate if these settings are also optimal for cyclic peptide-protein docking, the 50STR_FLEX protocol was performed in which peptides are defined as fully flexible during the it1 and itw stages of docking. Besides 50STR_FLEX, the 50STR_SOLVSHELL setup was tested in which complexes undergo explicit solvent shell refinement via a short MD simulation in water. Finally, the 50STR_FLEX_SOLVSHELL protocol was evaluated, including both full flexibility and explicit solvent refinement. The results were compared with the success rates of the 50STR protocol in which both extra flexibility and explicit solvent shell refinement were turned off (Table 1).

When considering the complete Backbone dataset, no clear answers are obtained regarding the impact of extra flexibility and explicit solvent-shell refinement on the quality of the generated models (Figure S4). However, as shown in Figure S5, increased flexibility does appear to generate more acceptable or higher quality models for the long peptides. Therefore, short and long peptides were analyzed separately to investigate the effect of peptide length on the prediction performance (Figure 6).

**Figure 6:**
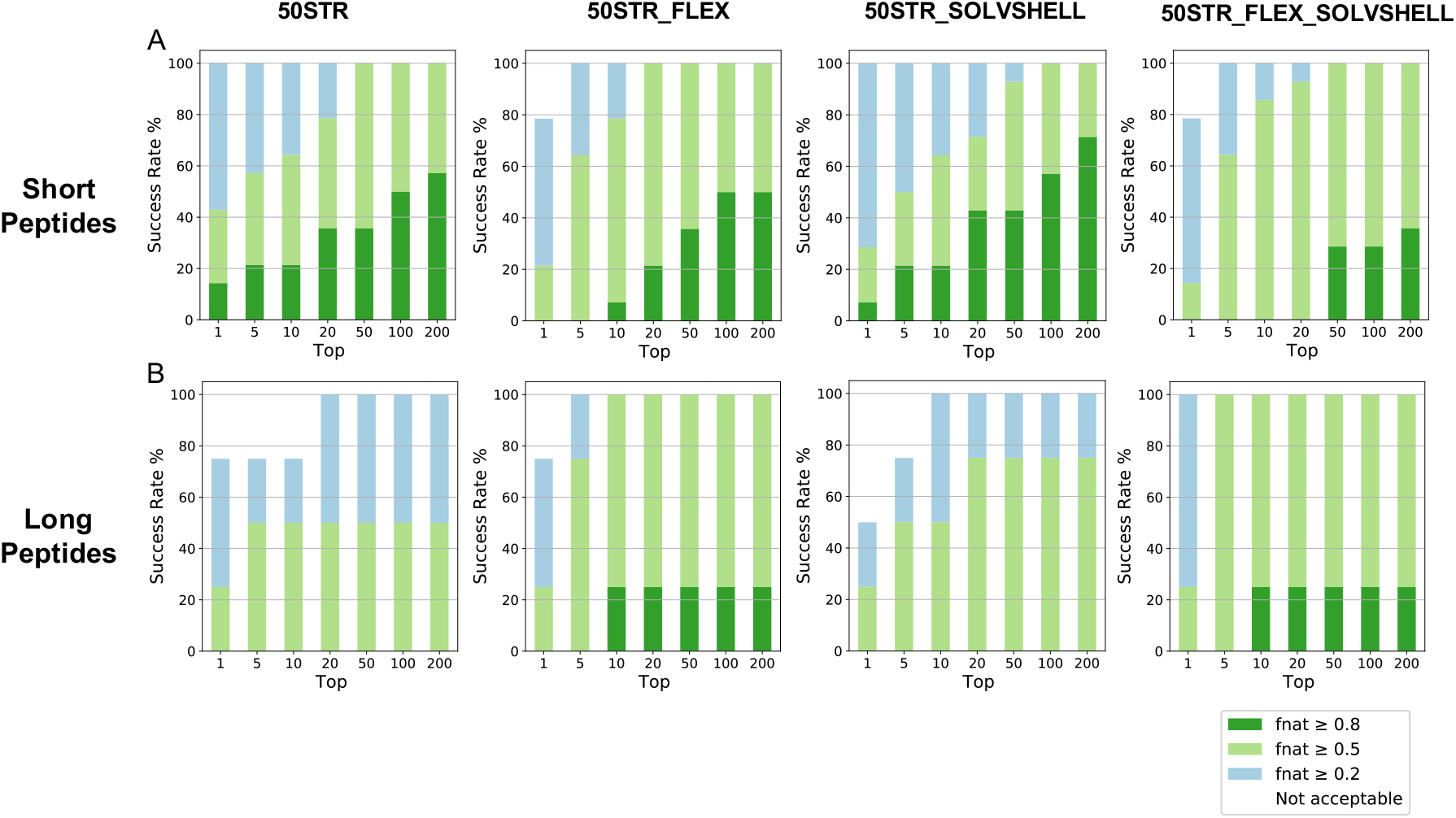
F_nat_ success-rate plots of 50STR, 50STR_SOLVSHELL and 50STR_FLEX docking protocols. (A) Plotted are the success rates for the short peptides (14 out of 18 complexes), ≤10 residues, of the Backbone dataset for four experiments (50STR, 50STR_FLEX, 50STR_SOLVSHELL and 50STR_FLEX_ SOLVSHELL). The color code (from blue to green) indicates model quality (from acceptable to high, respectively) according to CAPRI criteria. (B) Plotted are the success rates of the long peptides (4 out of 18 complexes), > 10 residues, of the dataset for four tested protocols (50STR, 50STR_FLEX, 50STR_SOLVSHELL and 50STR_FLEX_ SOLVSHELL). The color code (from blue to green) indicates model quality (from acceptable to high, respectively) according to CAPRI criteria.

For short cyclic peptides (≤10 residues), adding extra flexibility and/or explicit solvent refinement has no or a slightly negative impact on the docking performance (Figure 6A), whereas these two features do improve the prediction quality for long cyclic peptides (>10 residues) (Figure 6B), with the extra flexibility having the most impact. These results suggest cyclic peptides should be treated differently during docking according to their length. Long cyclic peptides (> 10 residues) show more similarities to linear peptides than short cyclic peptides and benefit from including extra flexibility and an explicit solvent shell. On the other hand, the optimal performance for short cyclic peptides (≤ 10 residues) is achieved by using the default flexibility setting as well as the default final energy minimization in itw (HADDOCK 2.4).

### The Optimized Docking Protocol Performs Better for the Backbone than for the Disulfide Dataset

After characterization of the optimal ensemble size and the effect of flexibility and solvent refinement on cyclic peptide docking, the best performing protocol was identified to be 50STR_COMB. This protocol uses a maximum of 50 structures in the peptide ensemble that is used as input for docking and treats peptides differently during the docking protocol, depending on their sequence length. For short peptides, ≤10 residues, the 50STR protocol is applied, while the long peptides, > 10 residues, are docked using the 50STR_FLEX_ SOLVSHELL setup, allowing for extra peptide flexibility. When using this optimized docking protocol for the holo receptor, HADDOCK could generate acceptable or higher-quality solutions for every complex of the dataset within the top 10 HADDOCK-scored structures (Figure S6, 7). When individually assessing the Backbone and Disulfide dataset, HADDOCK shows success rates of 83.3% and 50% for medium or higher quality structures within the top 10 models (*holo* receptor setup), respectively (Figure 7A).

**Figure 7:**
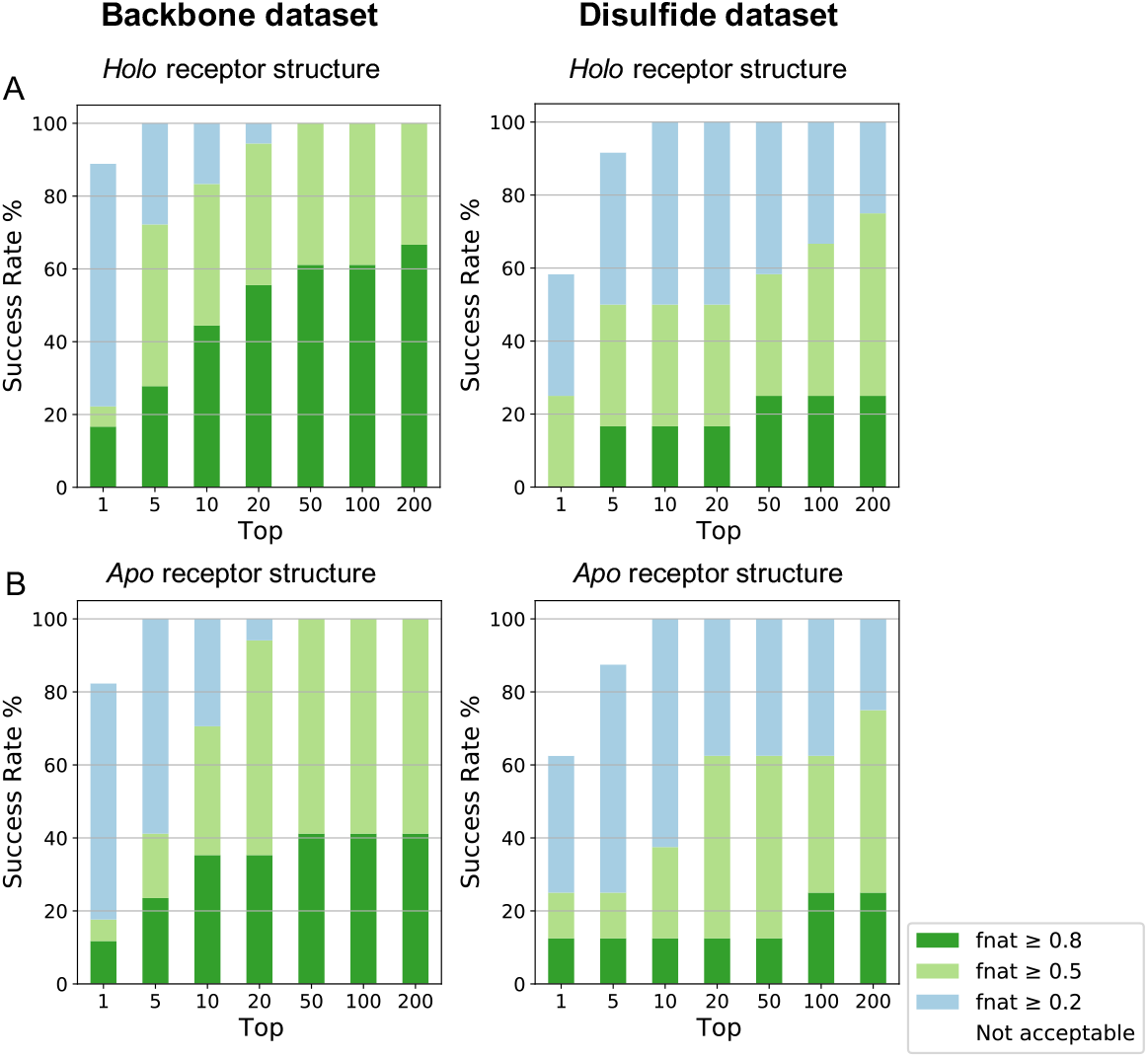
F_nat_ success-rate plots for 50STR_COMB docking protocol in both datasets. (A) Plotted are the docking success rates using the *holo* receptor for the Backbone (18 complexes) and the Disulfide dataset (12 complexes). Color coding (from blue to dark green) indicates model quality (from acceptable to high) according to CAPRI criteria. (B) Plotted are the docking success rates using the *apo* receptor for the Backbone (17 complexes) and the Disulfide dataset (8 complexes). Color coding (from blue to dark green) indicates model quality (from acceptable to high) according to CAPRI criteria.

The largest impact of introducing structural flexibility (it1 and itw) is on the model quality measured by f_nat_: From the it0 to the it1 stage, an f_nat_ improvement of almost 0.25 on average is obtained, while the effect of itw is less pronounced (0.05 on average) (Figure S7). This finding is in line with our previous work on peptide-protein docking^43^ showing that flexibility of the system is mainly handled during the flexible refinement stage (it1) of HADDOCK while the water refinement stage (itw) has the most impact on the energetics.

To determine the performance of HADDOCK in the more realistic apo setup (fully unbound), the optimal docking protocol (50STR_COMB) was performed using the *apo* form of the receptor and the generated peptide ensembles for both datasets. For the Backbone dataset, a success rate of 70.6% medium or higher quality in the top 10 solutions (Figure 7B) was obtained, while this was 37.5% for the Disulfide dataset (Figure 7B). Note that when considering acceptable or higher solutions, both holo and apo receptor docking lead to 100% success rate in the top 10 for both datasets, and even 100% in the top5 for the Backbone set.

The decrease in docking success between the Backbone and the Disulfide dataset mainly seems to be related to differences in the quality of the starting peptide conformational ensembles. As shown previously, the RMSD analysis of the peptide conformational ensembles revealed that the Backbone generated peptides were of better conformational quality than the Disulfide peptides due to a larger number of long peptides included in the Disulfide dataset with respect to the Backbone peptides (Figure 4). The docking performance difference between the two datasets appears to be related to the short/long peptide ratio of each dataset (Figure S5, S6, S8).

Analysis of the 50STR_COMB generated models showed that re-optimization of HADDOCK’s scoring function for cyclic peptides could be required to rank all high-to-medium quality models within the top 10. High-quality structures are predicted but these are often ranked among the top 100 solutions (Table S11), making it impossible for a user to extract them. Clustering of the models according to their interface ligand RMSD can partially compensate for the scoring challenges imposed as the model dimension is significantly reduced, leaving only representative models of each cluster to be considered by the user (Figure S2). These include the top 4 HADDOCK scored models per cluster which are used to calculate the overall cluster score (16 models for the best 4 clusters).

## Comparison with a Best-Case Scenario (Bound Docking)

The 50STR_COMB protocol has been determined as the optimal docking protocol for cyclic peptides and its performance in both the Backbone and the Disulfide dataset has been assessed. However, it is worthwhile comparing the success rates of the 50STR_COMB protocol with the success rates of the best case (but unrealistic) scenario in which the holo cyclic peptide conformation is used instead of a modelled 50-structures peptide ensemble to dock with the holo receptor. In this best case scenario, no conformational changes are required for an optimal interaction between peptide and receptor. Since the input of these bound docking experiments uses the holo X-ray structures, their success rates portray the optimal performance of HADDOCK given the knowledge of the binding site on the receptor and the absence of conformational changes (Figure S9).

The best scenario performance reaches 94.4% and 66.7% in the HADDOCK-scored top 10 for medium or higher quality models (83.3% and 41.6% for the top 1) in the Backbone and Disulfide datasets, respectively (Figure S9). In the 50STR_COMB protocol (Table 1, S6), the long peptides (> 10 residues) are defined as fully flexible and are refined in explicit solvent. The conformations can thus move away from their bound form during flexible refinement. On the other hand, only the default flexibility and energy minimization in itw are applied to the short peptides (≤10 residues), resulting in a reduction in conformational sampling for short peptides with respect to the long ones. When only the short peptides are considered, HADDOCK predicts their correct pose (medium-to-high quality structure) for 92.9% of the Backbone dataset and for 100% of the Disulfide dataset (Figure S10). These results indicate that HADDOCK is a powerful tool for short cyclic peptide-protein complex structure prediction when the peptide’s conformation is known (which could be the case, for example, when NMR data allow to define the conformation of the bound peptide). The comparison of these results with the presented unbound docking results (medium-to-high quality structure in 83.3%, Backbone, and 50%, Disulfide, of top 10 in Figure 7) demonstrates that the quality of the initial peptide structure(s) greatly determines the quality of the achievable docking results.

## Comparison with the State-of-the-art

### HADDOCK’s Cyclic Peptide-Protein Docking Performance is Comparable to AutoDock CrankPep

In Figure S11, the same dataset is used to compare the complex predictions of AutoDock CrankPep (ADCP) ^26^ with those obtained using our optimized protocol (50STR_COMB). The overall HADDOCK success rate, including both Backbone and Disulfide datasets, is 70% for medium quality structures or higher (top 10) when docked to the *holo* receptor, which is equal to the ADCP success rate. This drops to 60% when docked to the *apo* receptor, against 76% for ADCP.

When the cluster analysis is performed for the HADDOCK results, the success rate of the best 3 clusters is already competitive with ADCP, even in the challenging unbound docking scenario (Figure 8). HADDOCK’s performance in the best 3 and best 4 clusters becomes 66.7 % and 73.3% respectively in the *holo* condition and 64% and 76% for the *apo* setup (Figure 8).

**Figure 8:**
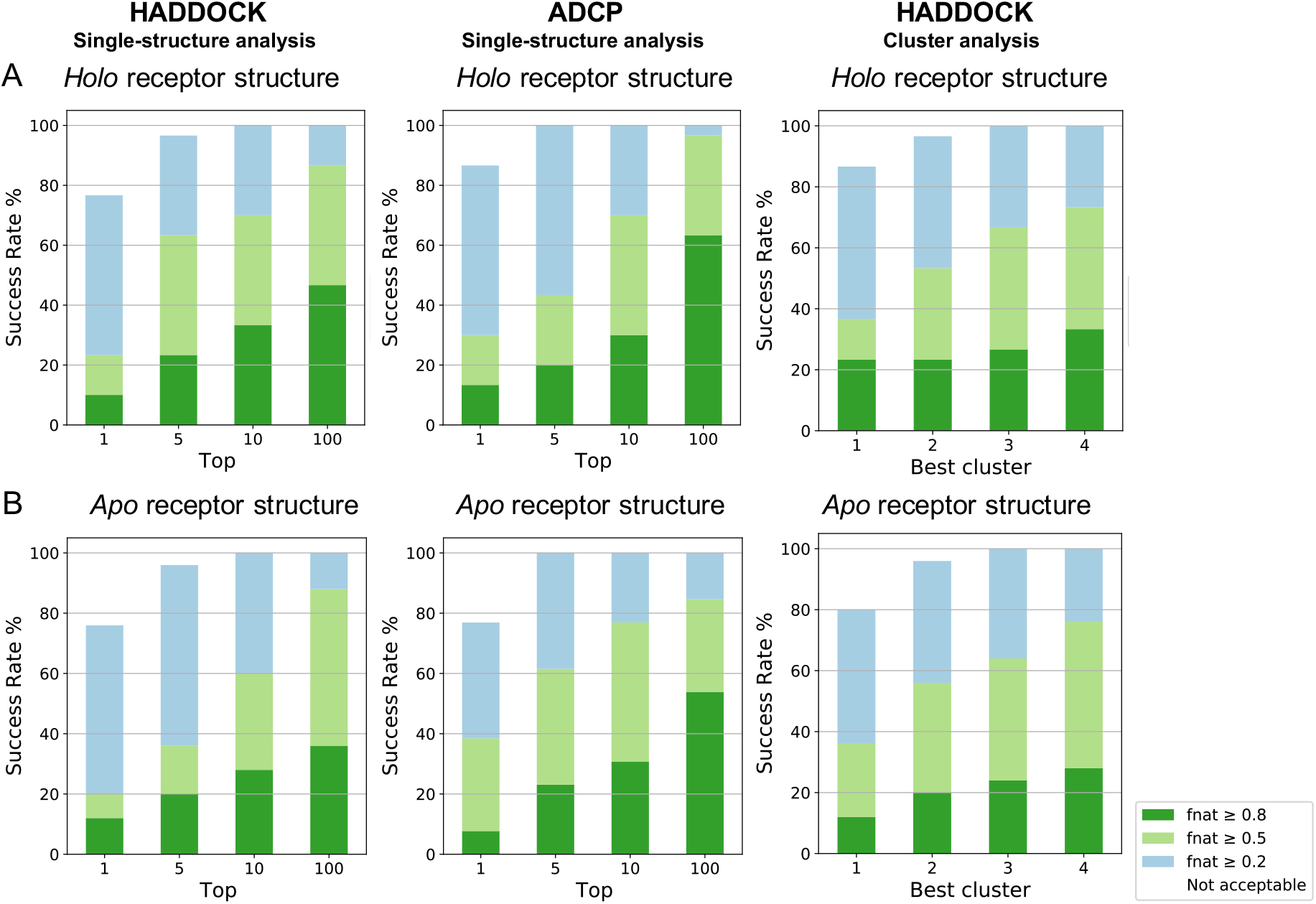
F_nat_ success-rate (%) plots to compare the performance of our optimized protocol with AutoDock CrankPep (ADCP). Plotted are the success rates of ADCP and HADDOCK, using either single-structure analysis or cluster analysis. (A) Plotted are the success rates from the *holo* receptor docking (30 complexes) and (B) the *apo* receptor runs (25 complexes). Color coding indicates model quality (from acceptable to high) according to CAPRI criteria.

When only the short peptides are considered (17 in total), the prediction’s performance improves compared to the complete dataset, reaching a success rate of 88.2% and 64.7% (top 10) for the *holo* and *apo* receptors, respectively. In the cluster analysis, the success rate of HADDOCK for medium or high-quality structures within the best 4 clusters becomes 94.1% using the *holo* receptor and 93.8% using the *apo*, a performance that is similar (or better in the *holo* receptor case) to ADCP. The performance for acceptable or better models is also in line with ADCP and even slightly better (1 complex difference) for the top 1 (Figure 9). Overall our results show that HADDOCK’s cyclic peptide-protein docking performance is similar to ADCP even in the challenging fully unbound docking condition. Performance is also comparable when considering only the clinically relevant^44^ short peptides (£10 residues), making HADDOCK a powerful tool for cyclic peptide-protein structure prediction.

**Figure 9:**
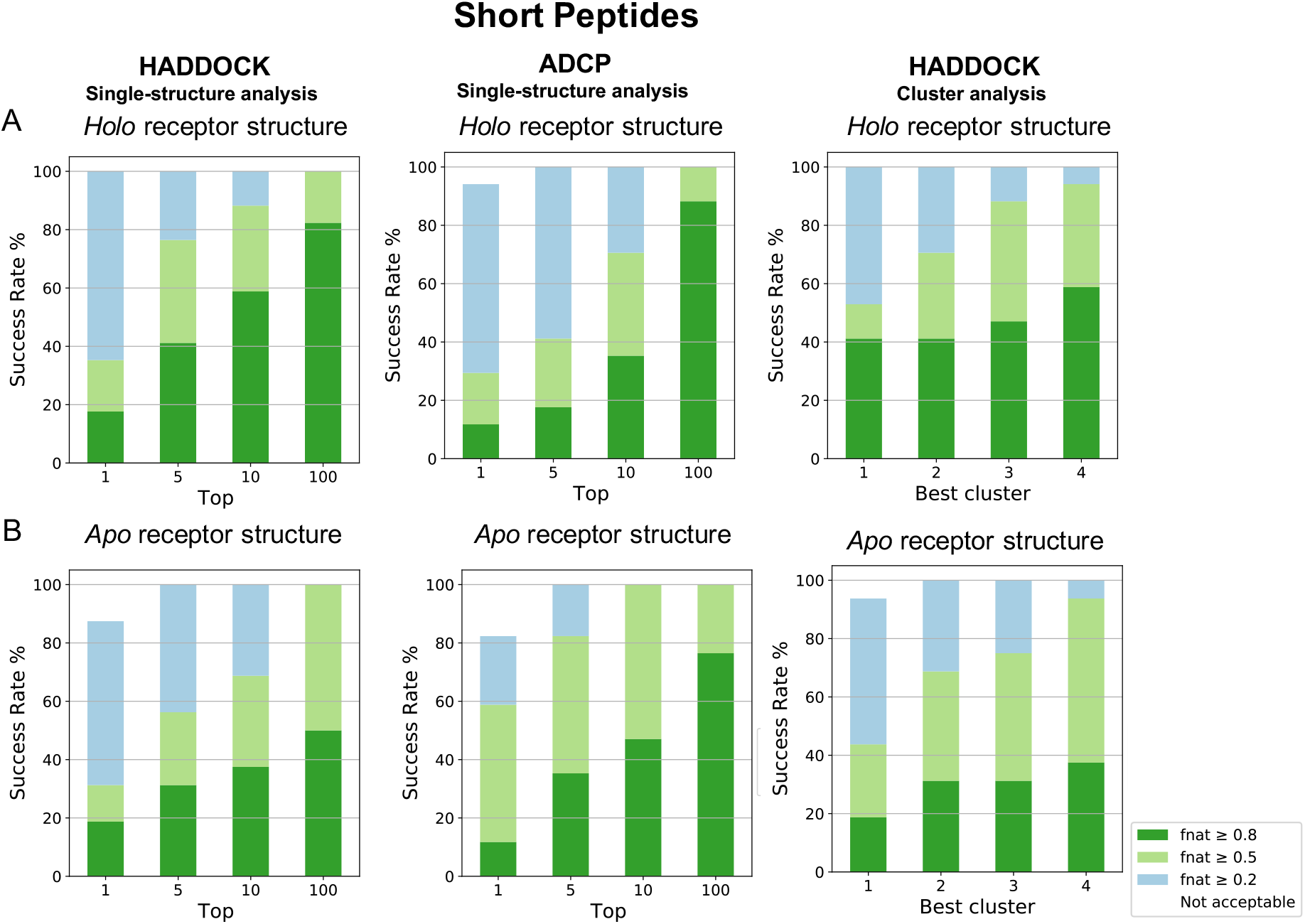
F_nat_ success-rate (%) plots for the short peptides in the two datasets. Plotted are the success rates of HADDOCK using single structure and cluster analysis and ADCP docking to the (A) holo (n=17) or (B) apo receptor structure (n=16). Color coding indicates model quality (from acceptable to high) according to CAPRI criteria.

## Conclusion

In this study, a step-by-step cyclisation and docking protocol based on the integrative modeling software HADDOCK2.4 has been described. By starting from the sequence of the cyclic peptides, conformational ensembles were generated, which were subsequently used for docking together with the respective *holo/apo* forms of the receptor. Analysis of the various protocols shows that peptides of different length should be treated differently, mainly to address the conformational flexibility challenge posed by longer peptides (>10 residues).

The ability of HADDOCK to predict cyclic peptide-protein complexes is comparable to the state-of-the-art practice in the field based on Autodock CrankPep (ADCP). Most importantly, HADDOCK performs similarly to ADCP in the case of fully unbound docking, and even slightly better when assessing the performance on a cluster basis (the default mode in HADDOCK). Since HADDOCK can also incorporate a variety of experimental data (e.g. NMR information about the peptide conformation) to guide the docking, the reported performance can be considered a lower limit, which can be enhanced by additional experimental input. In terms of computational efficiency, both ADCP and HADDOCK require a few hours or less to dock the peptides, depending on the number of CPUs available and the processor speed. For the peptide-protein docking with HADDOCK, users can also submit docking runs to the user-friendly webserver (https://wenmr.science.uu.nl/haddock2.4), which makes use of the European Open Science Cloud (EOSC) high throughput computing (HTC) resources^45^. Unfortunately, the generation of the cyclic peptide ensemble prior to docking is not yet supported by the webserver and should be run locally for the time being.

Finally, it is worth noting that the presented HADDOCK protocol performs particularly well in predicting short cyclic peptide-protein complexes (£10 residues), which constitute the majority of cyclic peptides in the clinic^44^. Thus, the presented cyclisation and docking protocol should be a valuable tool for rational design of clinically relevant cyclic peptides.

## Associated Content

### Supporting Information

The Supporting Information is available free of charge on the ACS Publications website at DOI:(DOI link) and includes:

- **Tables S1 and S2** Details of the peptide-protein complexes in the Backbone and the Disulfide dataset
- **Tables S3-S6** Parameters that have been changed in the run.cns and generate.inp files for Step 2 and Step 3 of cyclisation as well as for docking
- **Supplementary methods and Figure S1** Description of the difference between an f_nat_ and an i-RMSD analysis for the cyclic peptide-protein complexes
- **Tables S7-S10** RMSD analysis of the peptide conformational ensemble for the Backbone and the Disulfide dataset
- **Figure S2** Success-rate plots of the 50STR_COMB protocol, comparing single structure analysis and cluster analysis
- **Figure S3** Success-rate plots of the 60STR protocol
- **Figure S4** Success-rate plots of different docking protocols that all include a maximum of 50 peptide structures in the docking input
- **Figure S5 and S6** F_nat_ quality for each structure in the single structure analysis for different docking protocols that all include a maximum of 50 peptide structures in the docking input
- **Table S11** Structural quality analysis of the generated models from the optimized docking experiment (50STR_COMB)
- **Figure S9 and S10** Success-rate plots of the optimized protocol (50STR_COMB) for the Backbone and Disulfide dataset in the holo receptor (bound) condition
- **Figure S11** Success-rate plots of the optimized protocol (50STR_COMB) in comparison to ADCP’s performance

### Author

**Vicky Charitou** - *Bijvoet Center for Biomolecular Research, Faculty of Science - Chemistry, Utrecht University, Utrecht 3584CH, The Netherlands*

### Author Contributions

A.B. and S.K. designed and supervised the research with contributions from V.C. V.C and S.K. performed the benchmarking and data analysis. All the authors contributed to the writing of the manuscript.

### Funding

This work has been performed with the financial support of the Dutch Foundation for Scientific Research (NWO) (PPS Technology Area grant 741.018.201)) and the European Union Horizon 2020 project BioExcel (823830).

### Notes

The authors declare no competing financial interest.

The dataset used for this research and analysis scripts are available at https://github.com/haddocking/cyclic-peptides and the docking models are available from the SBGrid data reposity (https://data.sbgrid.org; DOI: …).

## Acknowledgements

The authors acknowledge all members from the Computational Structural Biology group at Utrecht University for fruitful discussions with a special mention of dr. Zuzana Jandová for her help with the peptide cyclisation protocol.

