## Supplementary Information for "A Cyclisation and Docking Protocol for Cyclic Peptide-Protein Modelling using HADDOCK2.4"

### Supplementary Methods

Another metric typically used for interface accuracy is the interface root-mean square deviation (i-RMSD), which is calculated on backbone atoms of both protein and peptide residues that are within a 10 Å radius from each other in the reference complex. However, i-RMSD fails to correctly evaluate acceptable cyclic peptide solutions as illustrated in Figure S1 in which the depicted model has the correct interface while only the flexible non-interacting peptide regions deviate from the crystal structure. This predicted complex scored high for $f_{nat}$ (0.86) with a i-RMSD higher than the acceptable cutoff (2.43Å), demonstrating that the i-RMSD metric can be misleading for the quality of cyclic peptide binding modes. This is because i-RMSD focuses on the backbone conformation of the peptide, while the crucial determinant for cyclic peptide interactions are the correct contacts, mainly obtained via side chains. The $f_{nat}$ metric is, therefore, ideal for reporting on the correct contacts and this is why it was chosen for the analysis in this study as well as in other studies concerning cyclic peptides[25].

*Table S1*. **The Backbone dataset including 18 complexes composed of a receptor and a peptide cyclized through its N- and C- terminus.** The length of the peptides varies from 6 to 14 residues and the corresponding *apo* structure of the receptor was found for all but one complex. The sequence of the peptides is shown with the additional disulfide bond highlighted in black in the respective cases, the termini are highlighted in boldface.

| PDB ID *holo receptor* | Peptide Length | PDB ID *apo receptor* [a] | Resolution (Å) | Sequence [b] | Identifier |
|---|---|---|---|---|---|
| **3wne** | 6 | 5jl4 | 1.70 | **P**KIDN**G** | 10.2210/pdb3WNE/pdb |
| **3zgc** | 7 | 1u6d | 2.20 | **G**DEETG**E** | 10.2210/pdb3ZGC/pdb |
| **3av9** | 8 | 5jl4 | 1.70 | **S**AKIDNL**D** | 10.2210/pdb3AV9/pdb |
| **3ava** | 8 | 5jl4 | 1.70 | **A**LKIDNL**D** | 10.2210/pdb3AVA/pdb |
| **3avb** | 8 | 5jl4 | 1.85 | **S**LKIDNL**D** | 10.2210/pdb3AVB/pdb |
| **3avf** | 8 | 5jl4 | 1.70 | **D**LKIDNL**D** | 10.2210/pdb3AVF/pdb |
| **3avg** | 8 | 5jl4 | 1.70 | **A**DKIDNL**D** | 10.2210/pdb3AVG/pdb |
| **3avh** | 8 | 5jl4 | 1.88 | **A**RKIDNL**D** | 10.2210/pdb3AVH/pdb |
| **3avi** | 8 | 5jl4 | 1.70 | **S**LKIDNM**D** | 10.2210/pdb3AVI/pdb |
| **3avj** | 8 | 5jl4 | 1.70 | **A**LKIDNM**D** | 10.2210/pdb3AVJ/pdb |
| **3avk** | 8 | 5jl4 | 1.75 | **S**LKIDNE**D** | 10.2210/pdb3AVK/pdb |
| **3avm** | 8 | 5jl4 | 1.88 | **S**RKIDNL**D** | 10.2210/pdb3AVM/pdb |
| **3avn** | 8 | 5jl4 | 2.10 | **S**HKIDNL**D** | 10.2210/pdb3AVN/pdb |
| **5xn3** | 8 | - | 1.34 | **R**GDINNN**V** | 10.2210/pdb5XN3/pdb |
| **1sfi** | 14 | 1tld | 1.65 | **G**RCTKSIPPICFP**D** | 10.2210/pdb1SFI/pdb |
| **3p8f** | 14 | 4is5 | 2.00 | **G**RCTKSIPPICFP**D** | 10.2210/pdb3P8F/pdb |
| **4k1e** | 14 | 4kga | 1.30 | **G**FCQRSIPPICFP**D** | 10.2210/pdb4K1E/pdb |
| **4kel** | 14 | 4kga | 1.15 | **G**FCQRSIPPICFP**N** | 10.2210/pdb4KEL/pdb |

a. Absence of the apo receptor structure is indicated with a dash ("- ")
b. Highlighted in black is the additional disulfide bond found in the peptides of the respective complexes. In black bold face the connection of the termini is represented.

*Table S2*. **Disulfide dataset including 12 complexes composed of a receptor and a peptide cyclized through a single disulfide bond.** The length of the peptides varies from 6-14 residues and the corresponding apo structure of the receptor was found for 8 out of 12 complexes. As shown in the sequence of each peptide by a black highlight, only one disulfide bond is present in each peptide that cyclizes the sequence.

| PDB ID *holo receptor* | Peptide Length | *apo* receptor PDB [a] | Resolution (Å) | Sequence [b] | Identifier |
|---|---|---|---|---|---|
| **3wnf** | 6 | 5jl4 | 1.45 | CKIDNC | 10.2210/pdb3WNF/pdb |
| **4ou3** | 6 | 4fkh | 1.95 | CNGRCG | 10.2210/pdb4OU3/pdb |
| **1smf** | 9 | 1tld | 2.10 | CTKSIPPEC | 10.2210/pdb1SMF/pdb |
| **3p72** | 11 | 1m0z | 1.90 | CTERMALHNLC | 10.2210/pdb3P72/pdb |
| **2ck0** | 11 | - | 2.20 | CKEWLSTAPCG | 10.2210/pdb2CK0/pdb |
| **5th2** | 12 | - | 1.84 | CQFDQSTRRLKC | 10.2210/pdb5TH2/pdb |
| **5djc** | 13 | 5dj0 | 2.10 | DCAWHLGELVWCT | 10.2210/pdb5DJC/pdb |
| **4ib5** | 13 | 3q9x | 2.20 | GCRLYGFKIHGCG | 10.2210/pdb4IB5/pdb |
| **5h5q** | 13 | 2obi | 1.10 | CRVDLQGWRRCRR | 10.2210/pdb5H5Q/pdb |
| **5eoc** | 13 | - | 1.98 | CQLINTNGSWHIC | 10.2210/pdb5EOC/pdb |
| **5wxr** | 14 | 4dva | 1.75 | GACSARGLENHAAC | 10.2210/pdb5WXR/pdb |
| **4m1d** | 14 | - | 1.80 | CRIHIGPGRAFYTC | 10.2210/pdb4M1D/pdb |

a. Absence of the apo receptor structure is indicated with a dash (“- “)
b. Highlighted in black is the disulfide bond that cyclizes the peptide.

*Table S3*. **Parameters changed in the `run.cns` file for Step 2 of the cyclisation protocol.** The indicated values are the optimized settings for Step 2 of cyclisation.

| Cyclisation Protocol: Step 2 | | | | |
|---|---|---|---|---|
| **Description** | **Keyword Changes** | | | |
| **Number of structures to dock** | structures_0=400 | structures_1=400 | waterrefine=400 | anastruc_1=400 |
| **Electrostatics** | elecflag_0=false | elecflag_1=false | | |
| **Explicit solvent refinement** | solvshell=true | | | |
| **Extra flexibility**[a] | nfle_1=1 | start_fle_1_1="1" | end_fle_1_1="14" | |
| **it0** | crossdock=false | randorien=false | rigidmini=false | ntrials=1 |
| **it1** | initiosteps=2000 | cool1_steps=2000 | cool2_steps=4000 | cool3_steps=4000 |
| **Clustering** | clust_meth="RMSD" | clust_cutoff=2.5 | | |

a. These parameters correspond to a cyclic peptide being defined as molecule 1 and consisting of 14 residues.

*Table S4*. **Parameters changed in the run.cns file for Step 3 of the cyclisation protocol.** Highlighted in black are the additional settings which are unique for Step 3 and not found in Step 2. The indicated values are the optimized settings for Step 3 of cyclisation.

| Cyclisation Protocol: Step 3 | | | | |
|---|---|---|---|---|
| **Description** | **Keyword Changes** | | | |
| **Number of structures to dock** | structures_0=400 | structures_1=400 | waterrefine=400 | anastruc_1=400 |
| **Electrostatics** | elecflag_0=false | elecflag_1=false | | |
| **Explicit solvent refinement** | solvshell=true | watersteps=5000 | | |
| **Extra flexibility**[a] | nfle_1=1 | start_fle_1_1="1" | end_fle_1_1="14" | |
| **it0** | crossdock=false | randorien=false | rigidmini=false | ntrials=1 |
| **it1** | initiosteps=2000 | cool1_steps=2000 | cool2_steps=4000 | cool3_steps=4000 |
| **Clustering** | clust_meth="RMSD" | clust_cutoff=2.5 | | |
| **Random seed** [b] | iniseed=3 | | | |
| **Unambiguous restraints** | unamb_lastit=1 | | | |
| **Cyclic peptide setting** [c] | cyclicpept_mol1=true | | | |

a. These parameters correspond to a cyclic peptide being defined as molecule 1 and consisting of 14 residues.

b. The value used as a random seed is different for every repetition of Step 3. In this example the random seed used is 3.

c. This parameter corresponds to a cyclic peptide being defined as molecule 1.

*Table S5*. **Parameters changed in `generate.inp` file (in the protocols directory of the run) for Step 3 of the cyclisation protocol.** Indicated values are the optimized settings for Step 3 of cyclisation.

| Cyclisation Protocol: Step 3 | |
|---|---|
| **Description** | **Keyword Changes** |
| **Minimum distance between C- and N-termini to create hydrogen bond** | `cyclicpept_dist=3.5` |
| **Minimum distance between cysteines to create a disulfide bond** | `disulphide_dist=4.0` |

*Table S6*. **Parameters changed in the `run.cns` file for all nine docking protocols tested in this study.** The indicated values are the optimized settings for the docking runs.

| Docking Protocol | | | | |
|---|---|---|---|---|
| **Description** | **Keyword Changes** | | | |
| **it1** | `initiosteps=2000` | `cool1_steps=2000` | `cool2_steps=4000` | `cool3_steps=4000` |
| **Number of structures to dock** | `structures_0=5000` | `structures_1=400` | `waterrefine=400` | `anastruc_1=400` |
| **Cyclic peptide setting [a]** | `cyclicpept_mol2=true` | | | |
| **Clustering** | `clust_meth="RMSD"` | `clust_cutoff=5` | | |
| **Hydrogen bond restraints** | `hbonds_on=true` | | | |

a. This parameter corresponds to a cyclic peptide being defined as molecule 2.

*Table S7*. **RMSD analysis of the Backbone peptide conformational ensemble.** Reported are the number of peptides within a given (high, medium and acceptable) quality. Peptides were defined as high quality if their RMSD (in relation to the x-ray structure) was lower than 1.5 Å, medium if RMSD lower than 2.0 Å and acceptable if their RMSD was lower than 2.5 Å. Complexes are sorted according to peptide's sequence length (from short to long peptides).

| Structure ensemble | Quality | PDB ID | | | | | | | | | | | | | | | | | |
|---|---|---|---|---|---|---|---|---|---|---|---|---|---|---|---|---|---|---|---|
| | | 3wne | 3zgc | 3av9 | 3ava | 3avb | 3avf | 3avg | 3avh | 3avi | 3avj | 3avk | 3avm | 3avn | 5xn3 | 4kel | 1sfi | 3p8f | 4k1e | Average |
| 5 | **High** | 1 | 1 | 0 | 0 | 0 | 0 | 0 | 0 | 0 | 0 | 0 | 0 | 0 | 0 | 0 | 0 | 0 | 0 | **0** |
| | **Medium** | 1 | 2 | 2 | 0 | 0 | 0 | 1 | 1 | 1 | 0 | 0 | 0 | 1 | 1 | 0 | 0 | 0 | 0 | **1** |
| | **Acceptable** | 1 | 2 | 2 | 1 | 1 | 1 | 1 | 1 | 1 | 0 | 0 | 0 | 1 | 1 | 0 | 0 | 1 | 0 | **1** |
| | **Total** | 2 | 5 | 5 | 5 | 5 | 5 | 5 | 5 | 5 | 5 | 5 | 5 | 5 | 5 | 5 | 5 | 5 | 5 | **5** |
| 30 | **High** | 3 | 1 | 0 | 0 | 0 | 1 | 2 | 1 | 0 | 2 | 0 | 0 | 0 | 1 | 0 | 0 | 0 | 0 | **1** |
| | **Medium** | 3 | 8 | 3 | 0 | 3 | 1 | 4 | 3 | 2 | 3 | 2 | 0 | 3 | 3 | 0 | 0 | 0 | 1 | **2** |
| | **Acceptable** | 4 | 10 | 6 | 2 | 5 | 7 | 6 | 6 | 7 | 10 | 4 | 0 | 4 | 3 | 0 | 0 | 3 | 1 | **4** |
| | **Total** | 8 | 21 | 30 | 30 | 29 | 30 | 30 | 30 | 30 | 30 | 30 | 30 | 30 | 23 | 30 | 30 | 30 | 30 | **28** |
| 40 | **High** | 4 | 2 | 0 | 0 | 0 | 1 | 4 | 2 | 0 | 2 | 0 | 0 | 0 | 1 | 0 | 0 | 0 | 0 | **1** |
| | **Medium** | 4 | 10 | 3 | 0 | 3 | 2 | 6 | 4 | 2 | 5 | 4 | 0 | 3 | 4 | 0 | 0 | 0 | 1 | **3** |
| | **Acceptable** | 5 | 12 | 8 | 2 | 7 | 8 | 8 | 8 | 10 | 14 | 6 | 1 | 4 | 4 | 0 | 0 | 3 | 3 | **6** |
| | **Total** | 10 | 26 | 39 | 40 | 39 | 40 | 40 | 40 | 40 | 40 | 40 | 40 | 40 | 32 | 40 | 40 | 40 | 40 | **37** |
| 50 | **High** | 5 | 3 | 1 | 0 | 1 | 1 | 4 | 2 | 0 | 3 | 0 | 1 | 0 | 2 | 0 | 0 | 0 | 0 | **1** |
| | **Medium** | 5 | 11 | 4 | 1 | 4 | 3 | 7 | 5 | 2 | 6 | 5 | 1 | 3 | 5 | 0 | 0 | 1 | 1 | **4** |
| | **Acceptable** | 6 | 13 | 10 | 3 | 8 | 10 | 9 | 9 | 10 | 17 | 7 | 2 | 7 | 6 | 1 | 1 | 4 | 3 | **7** |
| | **Total** | 12 | 33 | 49 | 50 | 49 | 50 | 50 | 50 | 50 | 50 | 50 | 50 | 50 | 42 | 50 | 50 | 50 | 50 | **46** |
| 60 | **High** | 6 | 5 | 1 | 0 | 1 | 1 | 4 | 3 | 0 | 3 | 0 | 1 | 0 | 2 | 0 | 0 | 0 | 0 | **2** |
| | **Medium** | 7 | 13 | 4 | 1 | 5 | 3 | 9 | 7 | 4 | 8 | 6 | 2 | 3 | 6 | 0 | 0 | 1 | 1 | **4** |
| | **Acceptable** | 7 | 15 | 12 | 3 | 9 | 11 | 11 | 11 | 13 | 20 | 8 | 3 | 8 | 8 | 2 | 1 | 5 | 4 | **8** |
| | **Total** | 14 | 41 | 59 | 60 | 59 | 60 | 60 | 60 | 60 | 60 | 60 | 60 | 60 | 49 | 60 | 60 | 60 | 60 | **56** |

*Table S8*. **RMSD analysis of the Disulfide peptide conformational ensemble.** Reported are the number of peptides within a given (high, medium and acceptable) quality. Peptides were defined as high quality if their RMSD (in relation to the x-ray structure) was lower than 1.5 Å, medium if RMSD lower than 2.0 Å and acceptable if their RMSD was lower than 2.5 Å. Complexes are sorted according to peptide's sequence length (from short to long peptides) and residues that are not part of the cyclic region have been excluded from the RMSD calculation.

| Structure ensemble | Quality | PDB ID | | | | | | | | | | | | |
|---|---|---|---|---|---|---|---|---|---|---|---|---|---|---|
| | | 3wnf | 4ou3 | 1smf | 3p72 | 2ck0 | 5th2 | 5djc | 4ib5 | 5h5q | 5eoc | 5wxr | 4m1d | Average |
| 5 | **High** | 0 | 2 | 0 | 0 | 0 | 0 | 0 | 0 | 0 | 0 | 0 | 0 | **0** |
| | **Medium** | 2 | 3 | 0 | 0 | 1 | 0 | 0 | 0 | 0 | 0 | 0 | 0 | **1** |
| | **Acceptable** | 4 | 4 | 1 | 0 | 1 | 0 | 0 | 0 | 0 | 0 | 0 | 0 | **1** |
| | **Total** | 5 | 4 | 5 | 5 | 5 | 5 | 2 | 5 | 1 | 5 | 5 | 4 | **4** |
| 30 | **High** | 2 | 3 | 0 | 0 | 0 | 0 | 0 | 0 | 0 | 0 | 0 | 0 | **0** |
| | **Medium** | 8 | 6 | 0 | 0 | 1 | 0 | 0 | 0 | 0 | 0 | 0 | 0 | **1** |
| | **Acceptable** | 15 | 10 | 2 | 0 | 5 | 0 | 0 | 0 | 0 | 0 | 1 | 0 | **3** |
| | **Total** | 23 | 10 | 27 | 30 | 30 | 26 | 4 | 19 | 8 | 30 | 30 | 17 | **21** |
| 40 | **High** | 3 | 3 | 0 | 0 | 0 | 0 | 0 | 0 | 0 | 0 | 0 | 0 | **1** |
| | **Medium** | 11 | 7 | 0 | 0 | 1 | 0 | 0 | 0 | 0 | 0 | 0 | 0 | **2** |
| | **Acceptable** | 18 | 12 | 3 | 0 | 7 | 0 | 0 | 1 | 0 | 0 | 1 | 0 | **4** |
| | **Total** | 30 | 13 | 36 | 40 | 40 | 33 | 5 | 24 | 11 | 40 | 40 | 19 | **28** |
| 50 | **High** | 4 | 3 | 0 | 0 | 0 | 0 | 0 | 0 | 0 | 0 | 0 | 0 | **1** |
| | **Medium** | 14 | 10 | 0 | 0 | 1 | 0 | 0 | 0 | 0 | 0 | 0 | 0 | **2** |
| | **Acceptable** | 23 | 15 | 3 | 0 | 9 | 0 | 0 | 1 | 0 | 0 | 1 | 0 | **4** |
| | **Total** | 37 | 16 | 45 | 50 | 50 | 41 | 8 | 30 | 14 | 50 | 50 | 25 | **35** |
| 60 | **High** | 5 | 3 | 0 | 0 | 0 | 0 | 0 | 0 | 0 | 0 | 0 | 0 | **1** |
| | **Medium** | 15 | 12 | 0 | 0 | 1 | 0 | 0 | 0 | 0 | 0 | 0 | 0 | **2** |
| | **Acceptable** | 26 | 17 | 3 | 0 | 10 | 0 | 0 | 2 | 0 | 0 | 1 | 0 | **5** |
| | **Total** | 43 | 19 | 54 | 60 | 60 | 51 | 9 | 35 | 16 | 60 | 60 | 30 | **41** |

*Table S9*. **RMSD analysis of the Backbone peptide conformational ensemble**. Reported are the best, worst and median RMSD values of every complex in all the different ensemble sizes. Complexes are sorted according to peptide's sequence length (from short to long peptides).

| Backbone Dataset | | | | | | | | | | | | | | | |
|---|---|---|---|---|---|---|---|---|---|---|---|---|---|---|---|
| | 5-structures RMSD | | | 30-structures RMSD | | | 40-structures RMSD | | | 50-structures RMSD | | | 60-structures RMSD | | |
| | Best | Worst | Median | Best | Worst | Median | Best | Worst | Median | Best | Worst | Median | Best | Worst | Median |
| **3wne** | 0.65 | 3.18 | 1.92 | 0.65 | 3.61 | 2.75 | 0.65 | 3.61 | 2.75 | 0.65 | 3.61 | 2.74 | 0.65 | 3.61 | 2.74 |
| **3zgc** | 1.35 | 3.67 | 2.81 | 1.35 | 4.39 | 2.78 | 1.06 | 4.39 | 2.79 | 1.06 | 4.39 | 2.86 | 1.06 | 4.39 | 2.88 |
| **3av9** | 1.71 | 4.47 | 2.67 | 1.71 | 4.47 | 3.18 | 1.71 | 4.92 | 3.23 | 1.14 | 4.92 | 3.23 | 1.14 | 4.92 | 3.23 |
| **3ava** | 2.16 | 4.55 | 3.82 | 2.01 | 5.16 | 3.53 | 2.01 | 5.16 | 3.39 | 1.65 | 5.16 | 3.4 | 1.65 | 5.16 | 3.4 |
| **3avb** | 2.12 | 4.46 | 3.34 | 1.62 | 4.48 | 3.34 | 1.62 | 4.48 | 3.2 | 1.44 | 4.52 | 3.22 | 1.44 | 4.54 | 3.23 |
| **3avf** | 2.42 | 4.69 | 3.77 | 1.34 | 4.69 | 3.26 | 1.34 | 4.95 | 3.26 | 1.34 | 4.95 | 3.33 | 1.34 | 5.3 | 3.37 |
| **3avg** | 1.87 | 3.94 | 3.47 | 0.72 | 4.6 | 3.1 | 0.72 | 4.6 | 3.09 | 0.72 | 4.6 | 3.11 | 0.72 | 4.6 | 3.12 |
| **3avh** | 1.63 | 5.15 | 3.54 | 1.24 | 5.3 | 3.3 | 1.24 | 5.3 | 3.3 | 1.24 | 5.3 | 3.29 | 1.24 | 5.3 | 3.3 |
| **3avi** | 1.98 | 4.7 | 3.68 | 1.95 | 4.77 | 3.35 | 1.95 | 4.77 | 3.45 | 1.95 | 4.77 | 3.45 | 1.73 | 4.86 | 3.52 |
| **3avj** | 2.55 | 4.88 | 2.82 | 1.14 | 4.88 | 2.8 | 1.14 | 4.88 | 2.8 | 1.14 | 4.88 | 2.8 | 1.14 | 4.88 | 2.8 |
| **3avk** | 2.56 | 4.64 | 3.83 | 1.67 | 4.7 | 3.42 | 1.67 | 4.7 | 3.47 | 1.67 | 4.7 | 3.42 | 1.67 | 4.7 | 3.42 |
| **3avm** | 2.57 | 4.2 | 3.7 | 2.57 | 5.16 | 3.55 | 2.49 | 5.16 | 3.55 | 1.5 | 5.16 | 3.54 | 1.5 | 5.16 | 3.55 |
| **3avn** | 1.67 | 4.42 | 3.18 | 1.67 | 5.13 | 3.39 | 1.67 | 5.13 | 3.25 | 1.67 | 5.13 | 3.25 | 1.67 | 5.29 | 3.38 |
| **5xn3** | 1.91 | 4.48 | 2.54 | 1.32 | 4.54 | 3.27 | 1.32 | 4.54 | 3.2 | 1.31 | 4.72 | 3.05 | 1.31 | 4.88 | 2.96 |
| **1sfi** | 2.96 | 4.57 | 3.9 | 2.73 | 5.19 | 3.89 | 2.73 | 5.63 | 3.94 | 2.15 | 5.63 | 3.94 | 2.15 | 5.63 | 3.91 |
| **4kel** | 2.63 | 4.06 | 3.82 | 2.54 | 5.23 | 4.04 | 2.54 | 5.48 | 4.0 | 2.33 | 5.48 | 3.91 | 2.33 | 5.48 | 3.91 |
| **3p8f** | 2.27 | 5.2 | 4.09 | 2.19 | 5.45 | 4.02 | 2.19 | 5.45 | 3.96 | 1.94 | 5.45 | 3.87 | 1.94 | 5.45 | 3.84 |
| **4k1e** | 2.68 | 4.44 | 3.76 | 1.83 | 5.0 | 4.01 | 1.83 | 5.11 | 4.01 | 1.83 | 5.26 | 3.99 | 1.83 | 5.26 | 4.04 |
| **Average** | **2.09** | **4.43** | **3.37** | **1.68** | **4.82** | **3.39** | **1.66** | **4.90** | **3.37** | **1.49** | **4.92** | **3.36** | **1.47** | **4.97** | **3.37** |

*Table S10*. **RMSD analysis of the Disulfide peptide conformational ensemble.** Reported are the best, worst and median RMSD values of every complex in all the different ensemble sizes. Complexes are sorted according to peptide's sequence length (from short to long peptides). Residues that are not part of the cyclic region have been excluded from the RMSD calculation.

| Disulfide Dataset | | | | | | | | | | | | | | | |
|---|---|---|---|---|---|---|---|---|---|---|---|---|---|---|---|
| | 5-structures RMSD | | | 30-structures RMSD | | | 40-structures RMSD | | | 50-structures RMSD | | | 60-structures RMSD | | |
| | **Best** | **Worst** | **Median** | **Best** | **Worst** | **Median** | **Best** | **Worst** | **Median** | **Best** | **Worst** | **Median** | **Best** | **Worst** | **Median** |
| **3wnf** | 1.61 | 3.3 | 2.31 | 1.02 | 3.34 | 2.28 | 1.02 | 3.53 | 2.29 | 0.96 | 3.85 | 2.31 | 0.58 | 3.85 | 2.32 |
| **4ou3** | 1.01 | 2.28 | 1.48 | 1.01 | 2.29 | 1.88 | 1.01 | 2.6 | 1.88 | 1.01 | 2.6 | 1.86 | 1.01 | 2.6 | 1.84 |
| **1smf** | 2.24 | 3.88 | 2.78 | 2.24 | 4.62 | 3.32 | 2.24 | 4.62 | 3.21 | 2.24 | 4.62 | 3.23 | 2.24 | 4.62 | 3.23 |
| **3p72** | 3.67 | 5.12 | 4.22 | 3.29 | 5.26 | 4.58 | 3.29 | 5.26 | 4.51 | 3.29 | 5.5 | 4.59 | 3.29 | 5.59 | 4.61 |
| **2ck0** | 1.76 | 4.13 | 3.27 | 1.76 | 4.8 | 3.32 | 1.76 | 4.98 | 3.36 | 1.76 | 4.98 | 3.36 | 1.76 | 4.98 | 3.38 |
| **5th2** | 4.97 | 6.01 | 5.56 | 4.71 | 6.06 | 5.61 | 4.71 | 6.06 | 5.67 | 4.71 | 6.06 | 5.57 | 4.71 | 6.45 | 5.57 |
| **5djc** | 4.85 | 5.5 | 5.2 | 4.85 | 5.7 | 5.52 | 4.85 | 5.92 | 5.53 | 4.33 | 6.33 | 5.52 | 4.33 | 6.33 | 5.53 |
| **4ib5** | 3.32 | 6.54 | 5.0 | 2.84 | 6.54 | 4.95 | 2.4 | 6.54 | 4.98 | 2.4 | 6.54 | 4.92 | 2.4 | 6.54 | 4.88 |
| **5h5q** | 5.57 | 5.57 | 5.57 | 2.72 | 5.57 | 4.00 | 2.72 | 6.03 | 4.01 | 2.72 | 6.04 | 3.97 | 2.72 | 7.03 | 3.97 |
| **5eoc** | 6.23 | 6.59 | 6.32 | 5.06 | 7.08 | 6.26 | 4.79 | 7.08 | 6.26 | 4.79 | 7.08 | 6.26 | 4.79 | 7.08 | 6.33 |
| **5wxr** | 2.56 | 4.1 | 3.77 | 2.4 | 4.81 | 3.75 | 2.4 | 4.81 | 3.75 | 2.4 | 4.81 | 3.63 | 2.4 | 5.46 | 3.62 |
| **4m1d** | 4.06 | 6.13 | 4.93 | 4.06 | 7.59 | 5.31 | 4.06 | 7.59 | 5.31 | 4.05 | 7.62 | 5.31 | 4.0 | 8.0 | 5.34 |
| **Average** | **3.49** | **4.93** | **4.20** | **3.00** | **5.31** | **4.23** | **2.94** | **5.42** | **4.23** | **2.86** | **5.58** | **4.21** | **2.84** | **5.73** | **4.22** |

*Table S11*. **Analysis of 50STR_COMB generated models.** Reported are the rank of the first acceptable model (according to $f_{nat}$ and CAPRI criteria) and its $f_{nat}$ and i-RMSD values respectively. Also, the rank of the best $f_{nat}$ value is reported along with the i-RMSD and $f_{nat}$ value for each specific model. Finally, reported is the total number of high, medium and acceptable models in every complex docking run according to their $f_{nat}$ value and the CAPRI criteria. Complexes in the table are sorted according to the sequence length of their peptide (from short to long).

| 50STR_COMB | | | | | | | | | |
|---|---|---|---|---|---|---|---|---|---|
| | **1$^{st}$ acceptable model** | | | **Best $f_{nat}$ model** | | | **Total number of models** | | |
| | **Rank** | **i-RMSD** | **$F_{nat}$** | **Rank** | **i-RMSD** | **$F_{nat}$** | **High** | **Medium** | **Acceptable** |
| **3wne** | 1 | 1.53 | 0.91 | 7 | 1.2 | 0.96 | 32 | 72 | 222 |
| **3wnf** | 1 | 1.06 | 0.77 | 40 | 0.59 | 0.90 | 22 | 214 | 152 |
| **4ou3** | 1 | 1.77 | 0.45 | 4 | 0.61 | 0.87 | 32 | 117 | 194 |
| **3zgc** | 1 | 1.90 | 0.49 | 30 | 0.82 | 0.70 | 0 | 18 | 198 |
| **3av9** | 1 | 3.02 | 0.21 | 17 | 1.34 | 0.89 | 7 | 84 | 227 |
| **3avi** | 1 | 2.69 | 0.45 | 19 | 1.07 | 0.90 | 7 | 28 | 306 |
| **3avb** | 1 | 2.43 | 0.82 | 24 | 1.33 | 0.89 | 13 | 42 | 221 |
| **3avh** | 1 | 2.98 | 0.27 | 8 | 2.16 | 0.89 | 14 | 46 | 202 |
| **3avn** | 1 | 2.93 | 0.35 | 6 | 2.53 | 0.81 | 1 | 41 | 183 |
| **5xn3** | 1 | 2.91 | 0.29 | 172 | 1.62 | 0.64 | 0 | 23 | 272 |
| **3avf** | 1 | 2.45 | 0.30 | 21 | 2.07 | 0.89 | 9 | 83 | 184 |
| **3avk** | 1 | 1.98 | 0.82 | 43 | 1.90 | 0.86 | 7 | 43 | 157 |
| **3avm** | 1 | 1.92 | 0.56 | 14 | 1.24 | 0.85 | 1 | 29 | 240 |
| **3avg** | 1 | 2.99 | 0.27 | 8 | 1.86 | 0.96 | 8 | 54 | 199 |
| **3avj** | 1 | 2.69 | 0.34 | 10 | 1.14 | 0.84 | 1 | 28 | 259 |
| **3ava** | 1 | 3.00 | 0.32 | 313 | 2.23 | 0.88 | 2 | 45 | 283 |
| **1smf** | 1 | 1.29 | 0.71 | 50 | 0.96 | 0.89 | 8 | 137 | 189 |
| **3p72** | 1 | 3.07 | 0.27 | 185 | 2.01 | 0.49 | 0 | 0 | 96 |
| **2ck0** | 3 | 2.55 | 0.65 | 52 | 1.97 | 0.71 | 0 | 19 | 159 |
| **5th2** | 1 | 4.03 | 0.34 | 105 | 1.93 | 0.57 | 0 | 2 | 121 |
| **5djc** | 1 | 2.45 | 0.60 | 1 | 2.45 | 0.60 | 0 | 2 | 150 |
| **4ib5** | 3 | 4.01 | 0.26 | 95 | 3.32 | 0.54 | 0 | 7 | 127 |
| **5h5q** | 1 | 5.16 | 0.33 | 214 | 2.60 | 0.46 | 0 | 0 | 146 |
| **5eoc** | 6 | 5.88 | 0.20 | 25 | 3.74 | 0.46 | 0 | 0 | 70 |
| **5wxr** | 2 | 1.85 | 0.71 | 12 | 1.75 | 0.77 | 0 | 61 | 71 |
| **4k1e** | 1 | 2.81 | 0.41 | 41 | 1.65 | 0.68 | 0 | 12 | 125 |
| **4m1d** | 2 | 5.00 | 0.31 | 177 | 3.05 | 0.77 | 0 | 17 | 116 |
| **3p8f** | 2 | 1.89 | 0.53 | 19 | 1.45 | 0.66 | 0 | 11 | 93 |
| **1sfi** | 2 | 2.49 | 0.63 | 4 | 1.24 | 0.74 | 0 | 19 | 120 |
| **4kel** | 1 | 4.84 | 0.29 | 26 | 1.99 | 0.75 | 0 | 10 | 105 |

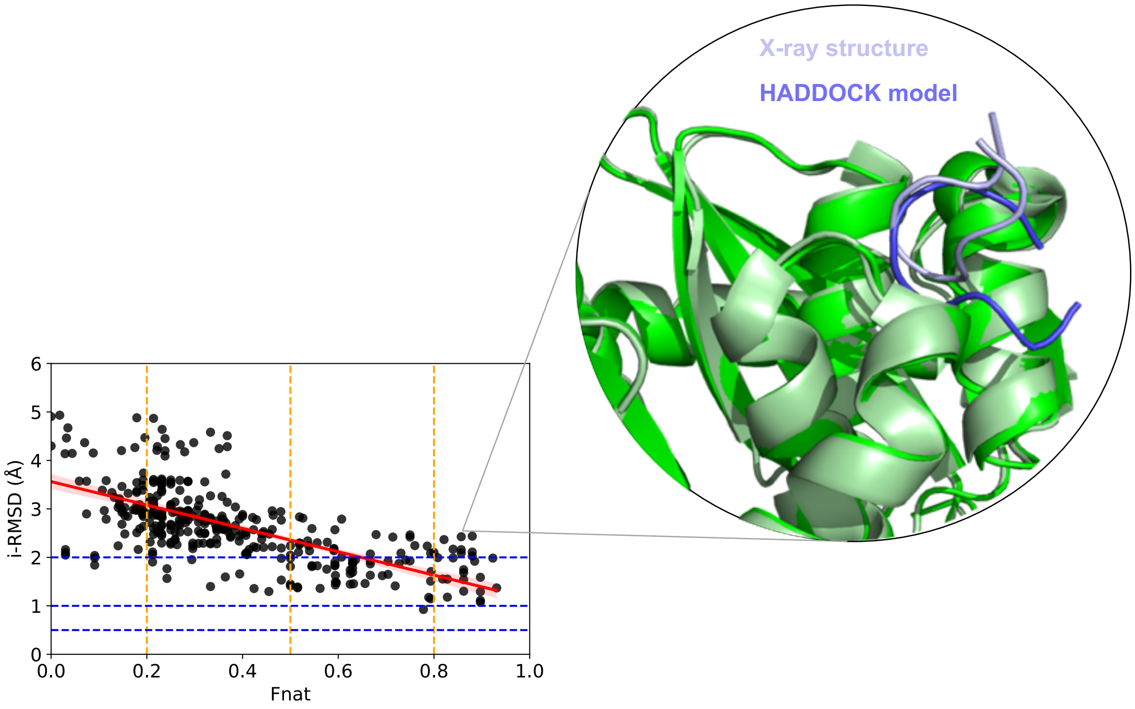

*Figure S1*. **Scatter plot of i-RMSD and $f_{nat}$ of top20 models for all complexes of the Backbone dataset, which were generated using the same protocol.** 360 data points were used to create the scatter plot. Illustrated in the circle is 3av9 complex model that scored high $f_{nat}$ (0.86) but also high i-RMSD (2.43).

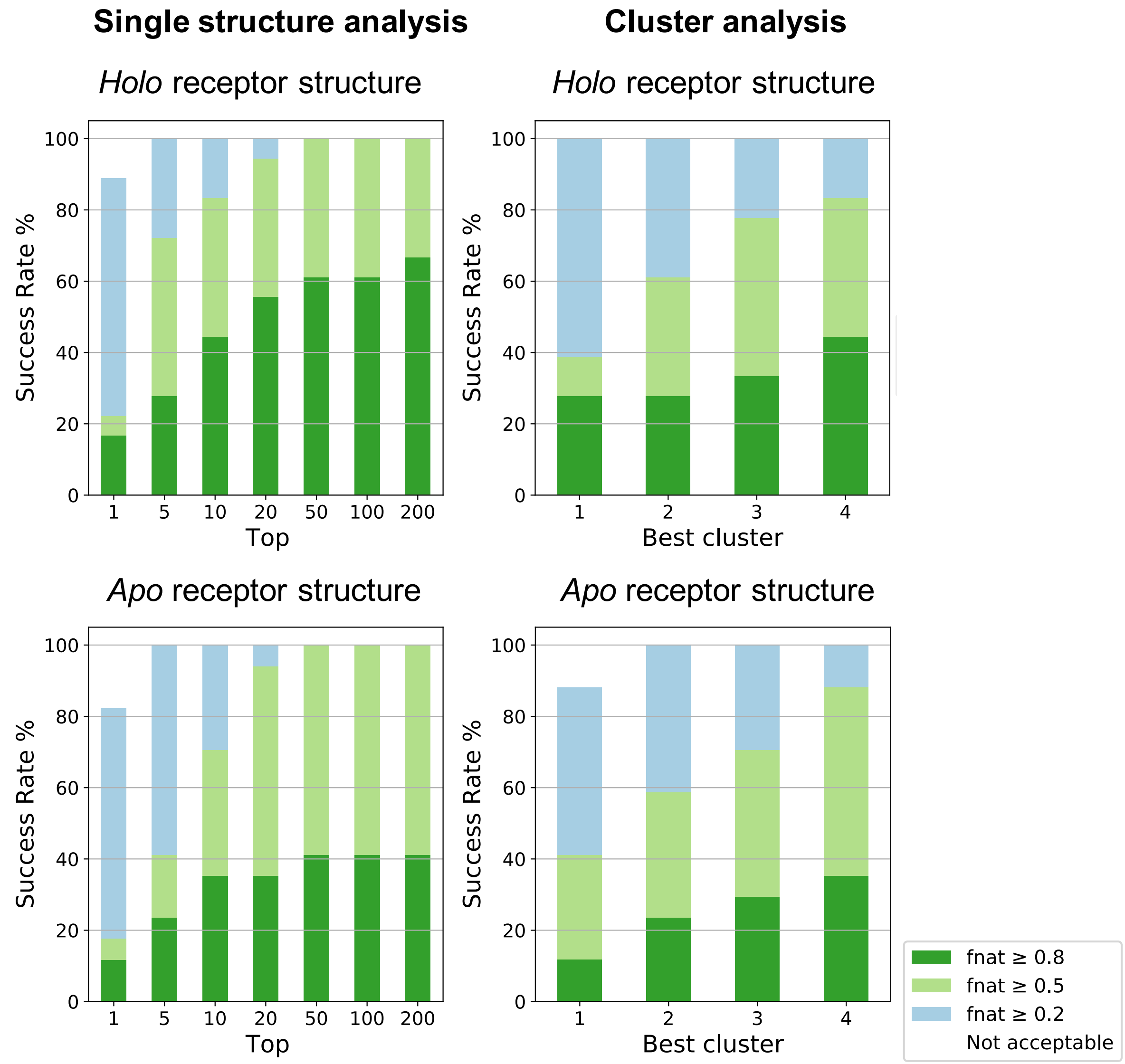

Figure S2. **$F_{nat}$ success rate (%) plots of the optimized experiment (50STR_COMB) for cyclic peptide using single structure or cluster analysis.** Plotted on the left are the success rates of the different tops (top 1, top 5, top 10 etc.) of the single structure analysis. On the right, accumulatively plotted are the success rates of the 4 best clusters, according to the HADDOCK score of itw. Color coding (from blue to green) indicates the quality of the models (from acceptable to high) according to CAPRI criteria.

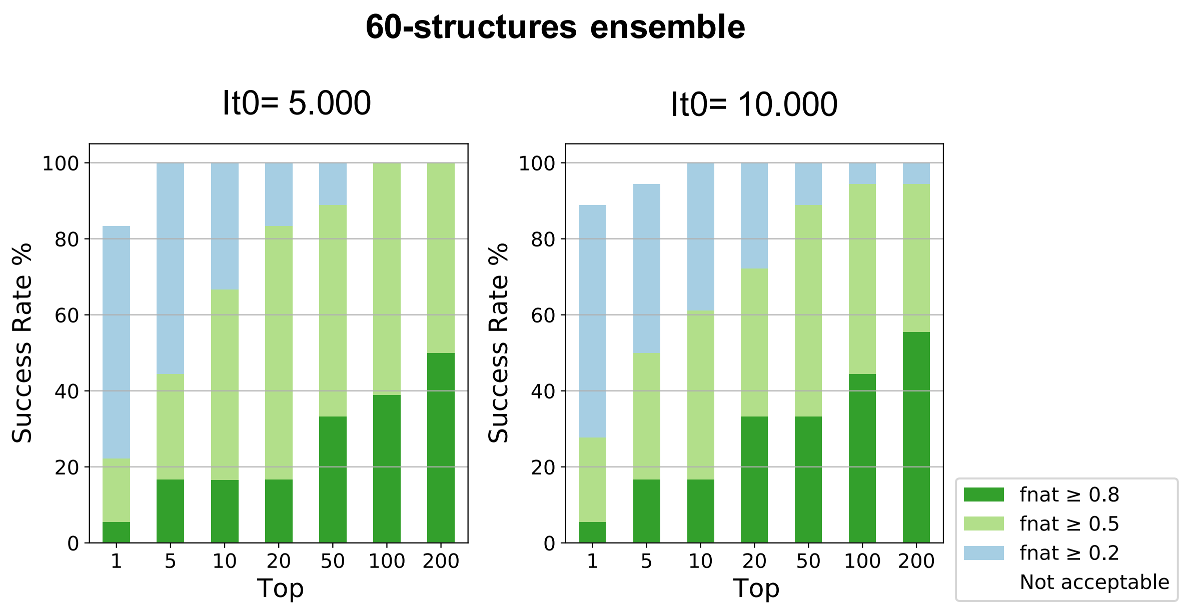

*Figure S3*. **$F_{nat}$ success rate (%) plots of experiments using a 60-structures ensemble and generating 5.000 or 10.000 models in it0.** Plotted are the success rates of the Backbone dataset (n=18 complexes). Color coding (from blue to green) indicates model quality (acceptable to high), according to CAPRI criteria.

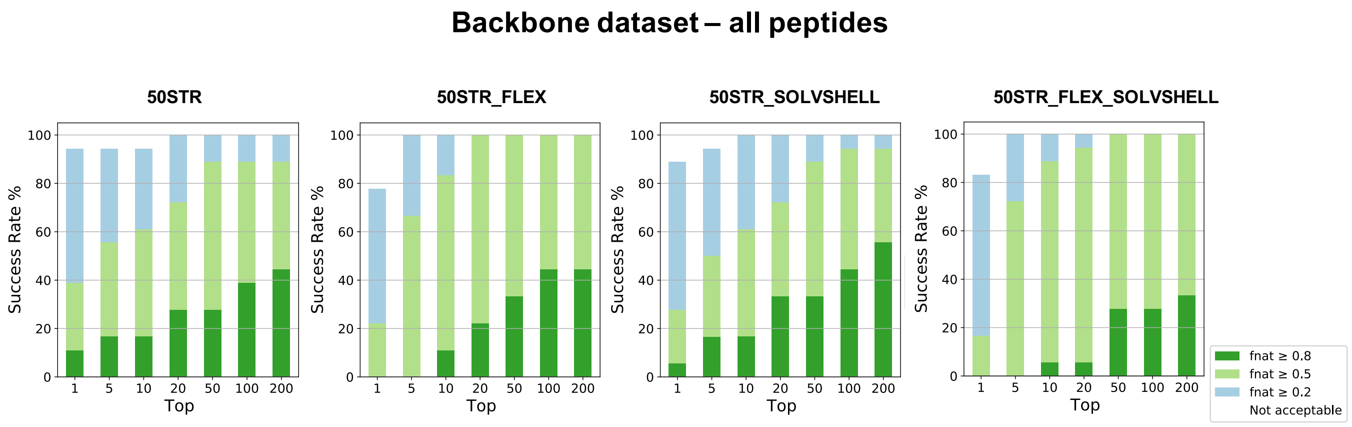

*Figure S4*. **$F_{nat}$ success-rate plots of 50STR_FLEX, 50STR_SOLVSHELL and 50STR docking protocols on the Backbone dataset.** Plotted are the success rates of all Backbone complexes for each tested protocol, the color coding (from blue to green) indicates the quality of the models (from acceptable to high) according to CAPRI criteria.

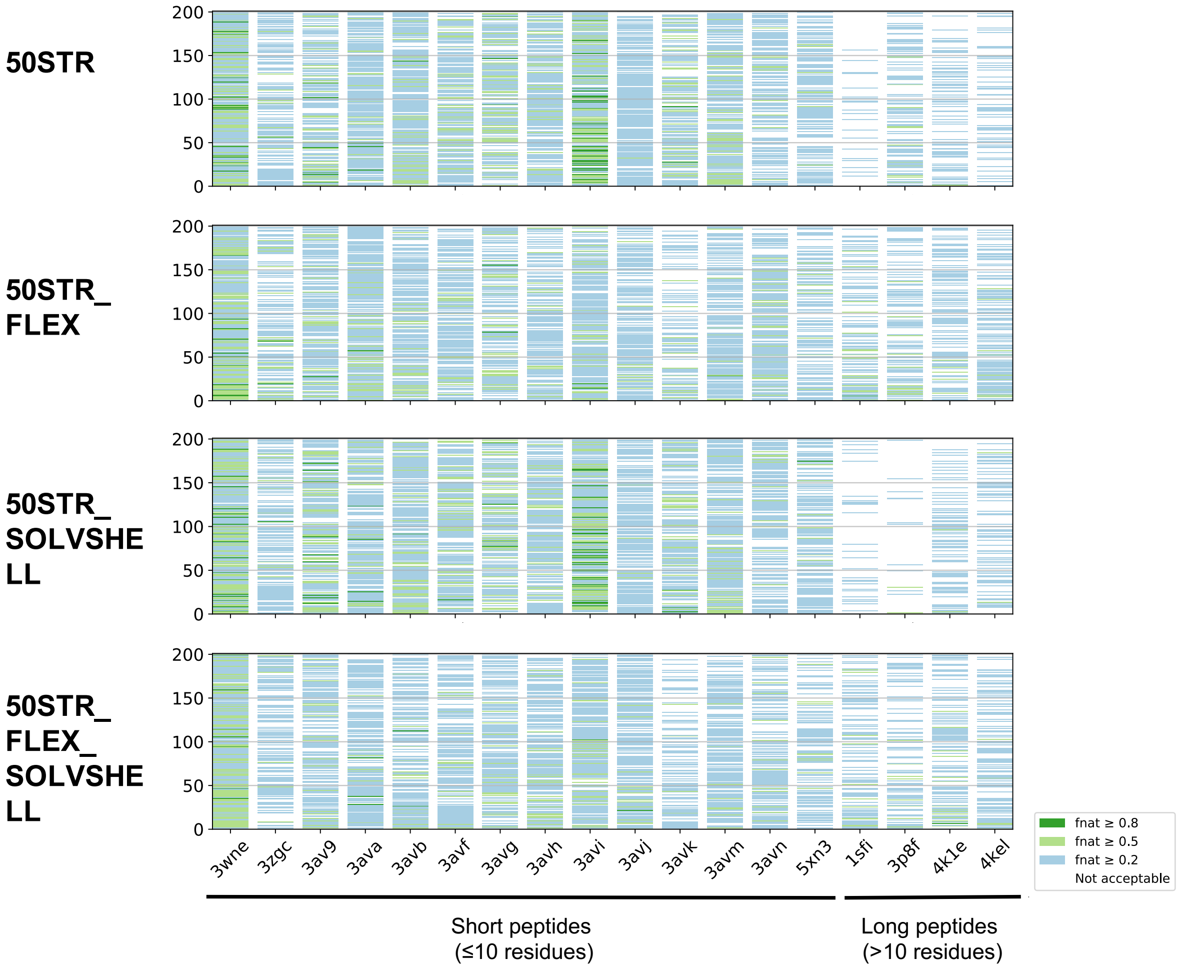

*Figure S5*. **Comparison of performance of 50STR, 50STR_FLEX, 50STR_SOLVSHELL and 50STR_FLEX_SOLVSHELL protocols.** Each column corresponds to one complex of the Backbone dataset and the y-axis reflects the ranking of the models according to the itw HADDOCK score in every protocol. The color of each model indicates its quality (from acceptable to high) according to the CAPRI criteria. Complexes are sorted in the x-axis according to their sequence length (from shorter to longer),

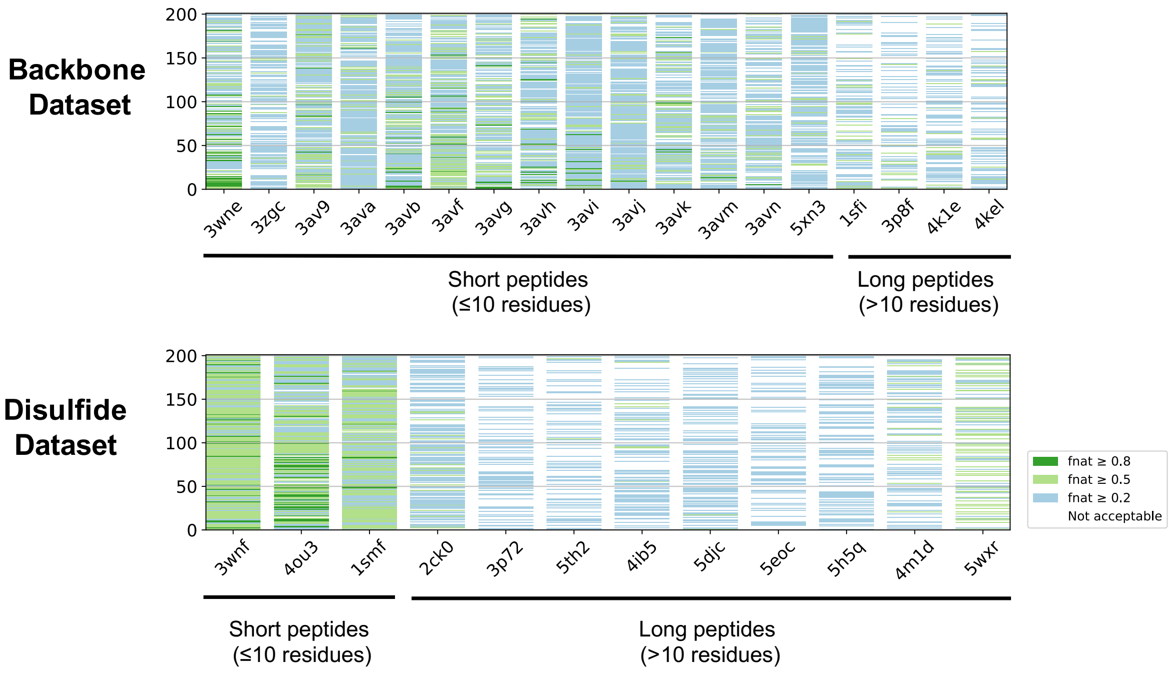

*Figure S6*. **Comparison of performance of 50STR_COMB experiment in the Backbone and Disulfide dataset.** Each column corresponds to one complex of the dataset and the y-axis reflects the ranking of the models according to the itw HADDOCK score in the protocol. The color of each model indicates its quality (from acceptable to high) according to the CAPRI criteria. Complexes are sorted in the x-axis according to their sequence length (from shorter to longer),

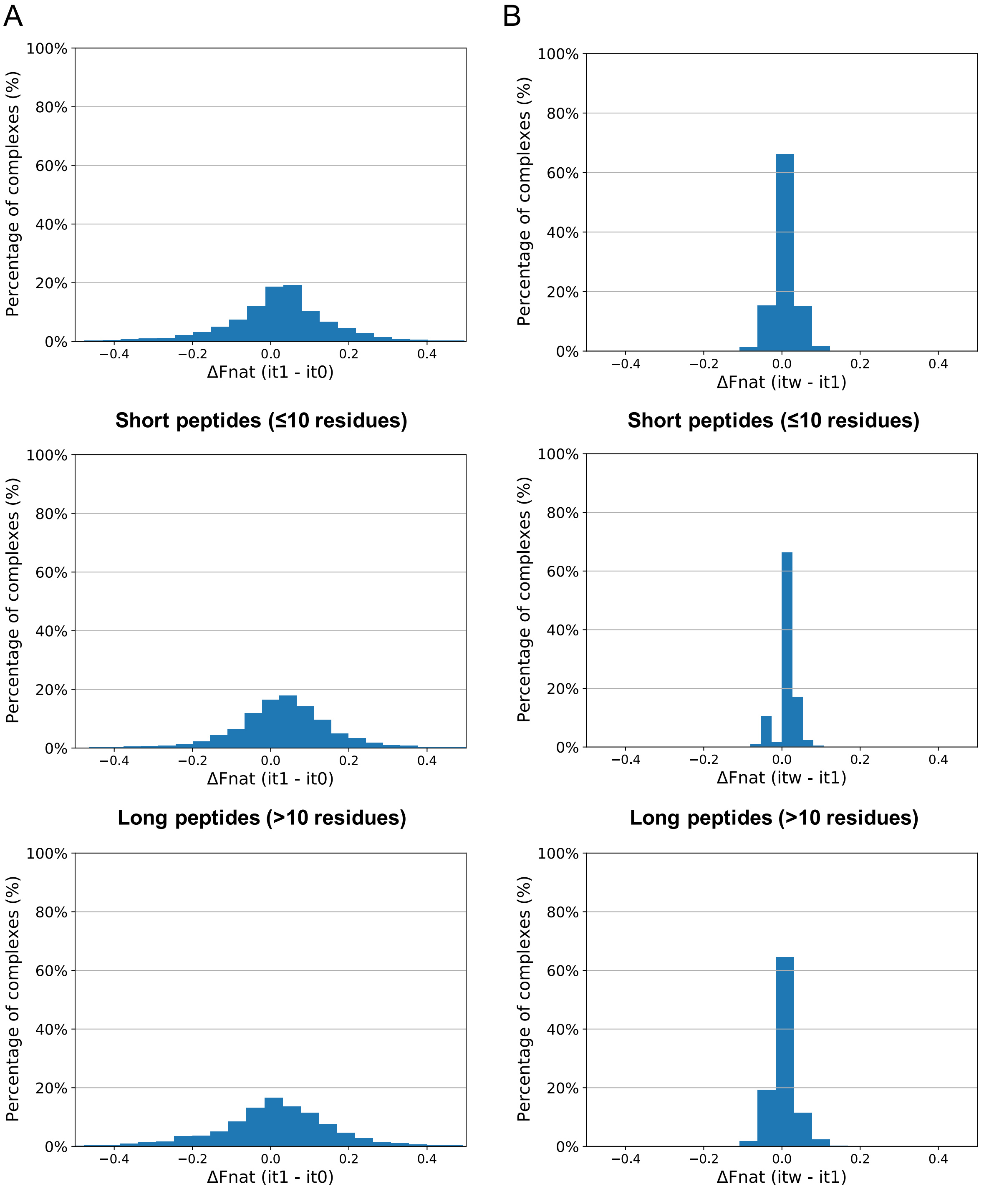

*Figure S7*. **Difference in $f_{nat}$ between models from various stages of HADDOCK (it0/it1/itw) for the 50STR_COMB experiment.** The distributions are calculated from all generated models of the 50STR_COMB experiment using the *holo* structure of the receptor (30 complexes*400 models/complex = 12.000 data-points). (A) The impact of semi-flexible refinement in torsion angle space (it1-it0) is shown. Bin size used for the plot: 35 (B) The impact of final water refinement (or the simple energy minimization step, if short peptides are considered) is shown. Bin size used for the plot: 10

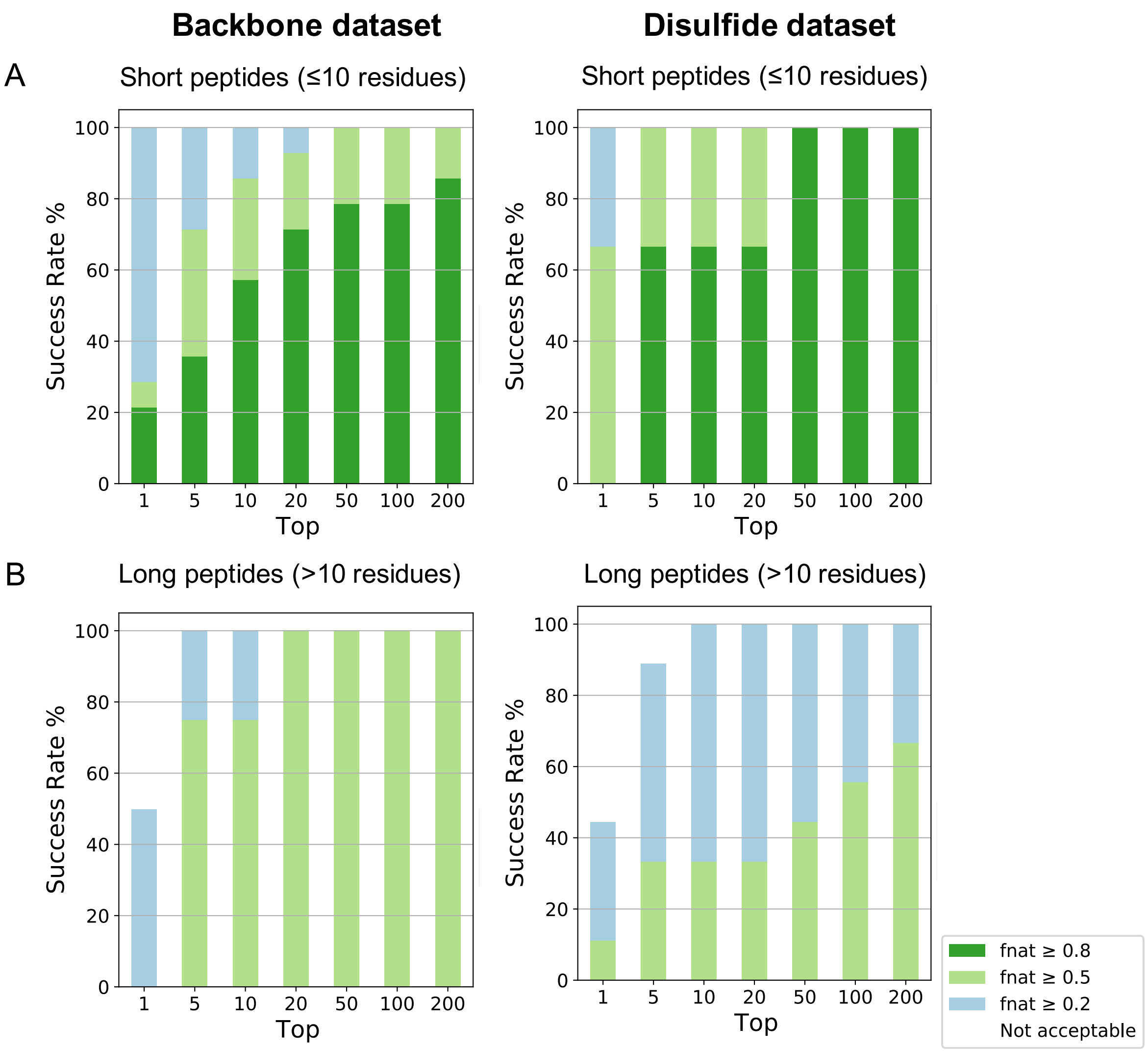

*Figure S8*. **$F_{nat}$ success-rate (%) plots of 50STR_COMB docking for short (≤10 residues) and long (>10 residues) peptides separately in the Backbone and Disulfide dataset.** (A) Plotted are the success-rates of the Backbone short peptides (n=14) and the Disulfide short peptides (n=3). (B) Plotted are the success-rates of the Backbone long peptides (n=4) and the Disulfide long peptides (n=9). Color coding (from blue to green) indicates the quality of models (from acceptable to high) according to CAPRI criteria.

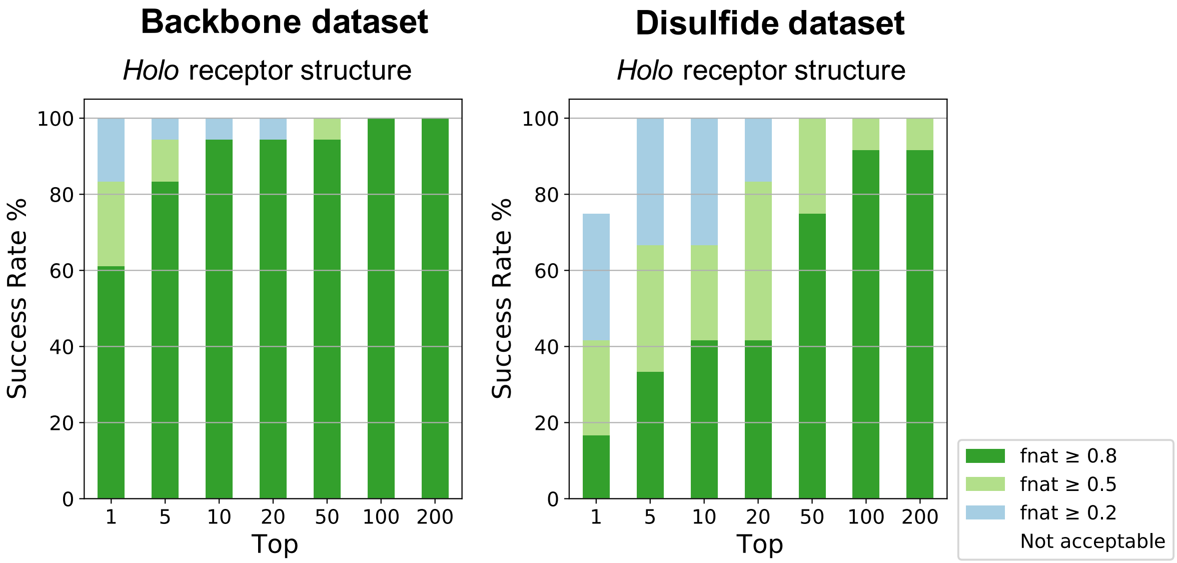

*Figure S9*. **$F_{nat}$ success-rate plots of best-case scenario (bound docking) for the Backbone and the Disulfide dataset.** Plotted are the success-rates of docking the *holo* cyclic peptide structure with the *holo* receptor structure for both datasets. Color coding (from blue to green) indicates model quality (from acceptable to high), according to CAPRI criteria.

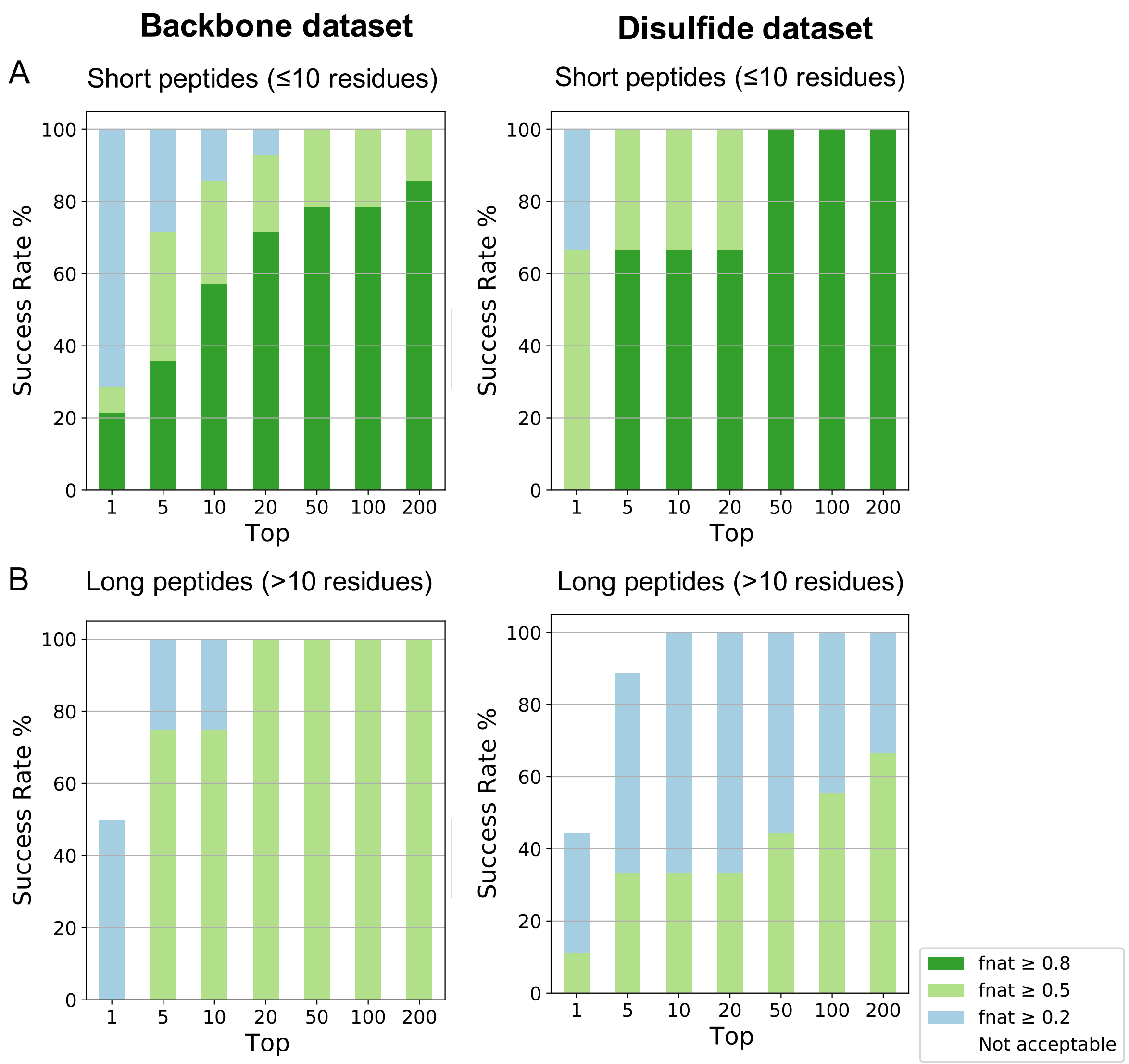

*Figure S10*. **$F_{nat}$ success-rate plots of best-case scenario (bound docking) for the Backbone and the Disulfide dataset.** (A) Plotted are the success-rates of short peptides using the *holo* receptor structure (B) Plotted are the success rates of long peptides using the *holo* receptor structure. Color coding (from blue to green) indicates model quality (from acceptable to high), according to CAPRI criteria.

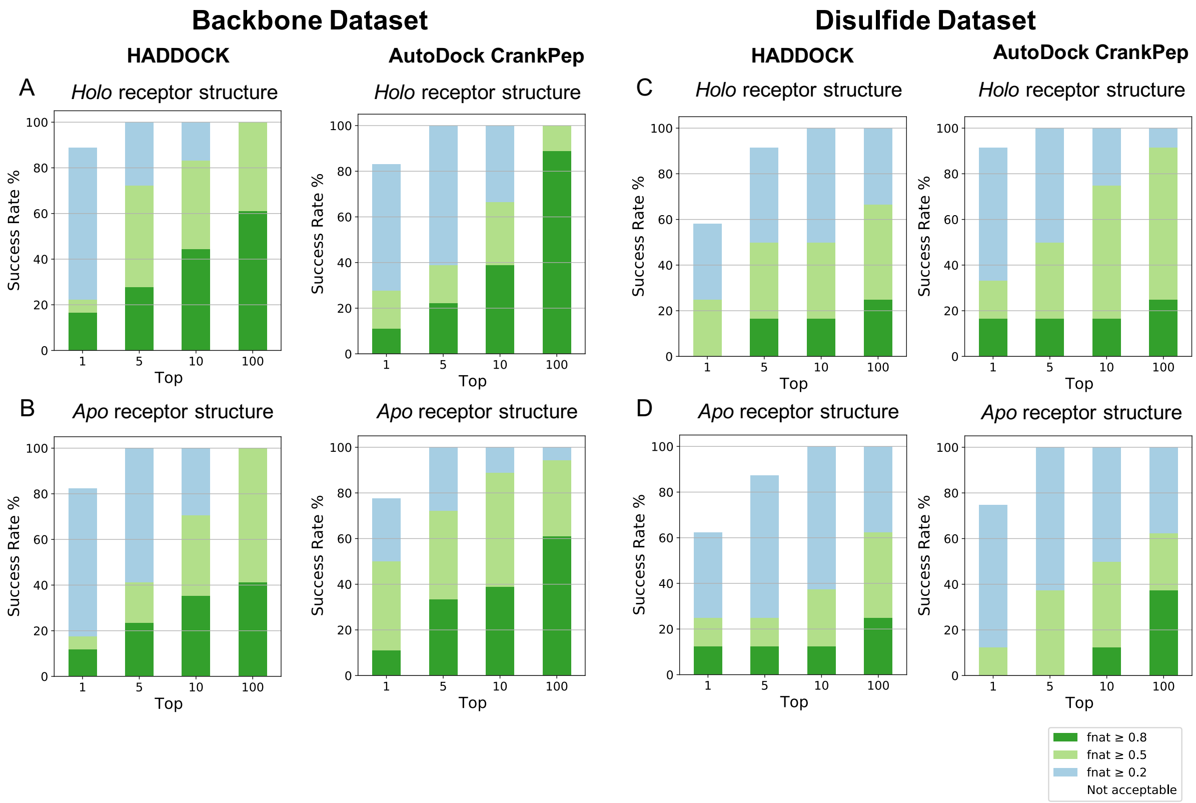

*Figure S11*. **Comparison of HADDOCK's and AutoDock CrankPep's performance.** (A) Plotted are the success-rates of the optimized HADDOCK protocol for cyclic peptides (50STR_COMB) and the ADCP protocol when using the Backbone dataset and the *holo* receptor (18 complexes) (B) when using the *apo* receptor (17 complexes) (C) Plotted are the success-rates of the HADDOCK protocol (50STR_COMB) and the ADCP protocol when using the Disulfide dataset and the *holo* receptor (12 complexes) (D) when using the *apo* receptor (8 complexes)
